# Upgraded molecular models of the human KCNQ1 potassium channel

**DOI:** 10.1101/648634

**Authors:** Georg Kuenze, Amanda M. Duran, Hope Woods, Kathryn R. Brewer, Eli Fritz McDonald, Carlos G. Vanoye, Alfred L. George, Charles R. Sanders, Jens Meiler

## Abstract

The voltage-gated potassium channel KCNQ1 (K_V_7.1) assembles with the KCNE1 accessory protein to generate the slow delayed rectifier current, I_KS_, which is critical for membrane repolarization as part of the cardiac action potential. Loss-of-function (LOF) mutations in KCNQ1 are the most common cause of congenital long QT syndrome (LQTS), type 1 LQTS, an inherited genetic predisposition to cardiac arrhythmia and sudden cardiac death. A detailed structural understanding of KCNQ1 is needed to elucidate the molecular basis for KCNQ1 LOF in disease and to enable structure-guided design of new anti-arrhythmic drugs. In this work, advanced structural models of human KCNQ1 in the resting/closed and activated/open states were developed by Rosetta homology modeling guided by newly available experimentally-based templates: *X. leavis* KCNQ1 and resting voltage sensor structures. Using molecular dynamics (MD) simulations, the models’ capability to describe experimentally established channel properties including state-dependent voltage sensor gating charge interactions and pore conformations, PIP2 binding sites, and voltage sensor – pore domain interactions were validated. Rosetta energy calculations were applied to assess the models’ utility in interpreting mutation-evoked KCNQ1 dysfunction by predicting the change in protein thermodynamic stability for 50 characterized KCNQ1 variants with mutations located in the voltage-sensing domain. Energetic destabilization was successfully predicted for folding-defective KCNQ1 LOF mutants whereas wild type-like mutants had no significant energetic frustrations, which supports growing evidence that mutation-induced protein destabilization is an especially common cause of KCNQ1 dysfunction. The new KCNQ1 Rosetta models provide helpful tools in the study of the structural mechanisms of KCNQ1 function and can be used to generate structure-based hypotheses to explain KCNQ1 dysfunction.

**Author Summary:** Cardiac rhythm is maintained by synchronized electrical impulses conducted throughout the heart. The potassium ion channel KCNQ1 is important for the repolarization phase of the cardiac action potential that underlies these electrical impulses. Heritable mutations in KCNQ1 can lead to channel loss-of-function (LOF) and predisposition to a life-threatening cardiac arrhythmia. Knowledge of the three-dimensional structure of KCNQ1 is important to understand how mutations lead to LOF and to support structurally-guided design of new anti-arrhythmic drugs. In this work, we present the development and validation of molecular models of human KCNQ1 inferred by homology from the structure of frog KCNQ1. Models were developed for the open channel state in which potassium ions can pass through the channel and the closed state in which the channel is not conductive. Using molecular dynamics simulations, interactions in the voltage-sensing and pore domain of KCNQ1 and with the membrane lipid PIP2 were analyzed. Energy calculations for KCNQ1 mutations in the voltage-sensing domain reveled that most of the mutations that lead to LOF cause energetic destabilization of the KCNQ1 protein. The results support both the utility of the new models and growing evidence that mutation-induced protein destabilization is a common cause of KCNQ1 dysfunction.

## Introduction

Voltage-gated ion channels are ubiquitously expressed in human tissues and contribute to diverse physiological phenomena such as generation and modulation of the membrane potential in excitable cells, myocyte contraction, modulation of neurotransmitter and hormone release, and electrolyte transport in epithelia. KCNQ1 (K_V_7.1) is a voltage-gated potassium (K_V_) channel expressed in the heart and in epithelial cells in the inner ear, stomach, kidney and colon (1). Like other K_V_ channels KCNQ1 is comprised of four identical α-subunits, each containing six membrane-spanning segments (S1-S6) and a pore loop (P loop) that contributes to the ion selectivity filter in the homo-tetramers to create the KCNQ1 channel. The central pore domain (PD, S5-P-S6) is surrounded by four voltage-sensing domains (VSDs, S1-S4) that respond to membrane depolarization by a conformational change that triggers structural rearrangements in the PD (called electromechanical coupling), which opens the channel gate making the channel conductive (2, 3). VSD activation occurs stepwise and proceeds from an initial resting VSD conformation in which the PD is closed (resting/closed, RC) to an activated VSD with an open PD (activated/open, AO) (4, 5). In KCNQ1, these transitions also involve an experimentally resolvable intermediate state (6, 7). A hallmark of the KCNQ1 channel is its co-assembly with the KCNE1 auxiliary subunit in the heart to generate the channel complex that is responsible for the slow delayed rectifier current (I_KS_) necessary for myocardial repolarization (8, 9).

Heritable mutations in KCNQ1 are associated with several cardiac diseases including long QT syndrome, atrial fibrillation, and short QT syndrome (10). About 50% of the genetic cases of long QT syndrome (LQTS), which predispose children and young adults to sudden cardiac death, are associated with dominant mutations in KCNQ1 (type 1 LQTS) (11). While progress in the functional characterization of LQTS-associated mutations has been made (12–15), the molecular mechanisms underlying channel dysfunction remain difficult to assess without the availability of high-accuracy structural data. A detailed molecular understanding is needed to improve decision making for new unclassified KCNQ1 mutations and to support the development of new anti-arrhythmic therapeutics.

The lack of an experimentally determined structure for human KCNQ1 has prompted molecular modeling efforts. The first structural model of the KCNQ1 channel containing all membrane-embedded regions S1-S6 was published in 2007 (16) and since then has stimulated numerous structural and functional studies that provided new insights into KCNQ1, e.g. its regulation by KCNE1 (17–20) and other KCNE proteins (21, 22), binding of phosphatidyl-4,5-bisphosphate (PIP2) (23, 24), the structural determinants and transitions in VSD activation (5, 25), and mechanisms of VSD-PD electromechanical coupling (26). The recent determination of a cryo-EM structure of X. *leavis* KCNQ1 (27) which shares 78% sequence identity to human KCNQ1 provides a new template to upgrade the first Rosetta homology models of KCNQ1. The PD of this structure is believed to be in its closed state conformation, while its VSD occupies the activated state. In addition to this cryo-EM structure, recent experimentally determined structures of non-mammalian VSDs in resting conformations (28, 29) provide new templates for modeling the resting/closed channel which previously was informed by purely computational models built by combining the bacterial KcsA template with *de novo* prediction methods (16, 30).

Here, using this new set of experimental structures leveraged in Rosetta multiple-template homology modeling, we report the development and structural validation of a second generation of Rosetta models of the human KCNQ1 channel in the closed and open states. These models have improved completeness and accuracy and thus represent useful tools for studying the structure-function relationships in KCNQ1 and the molecular mechanisms of mutation-induced changes in KCNQ1 phenotypes. We showcase the utility of the new models by (1) applying molecular dynamics (MD) simulation to our KCNQ1 models which generated useful insight into the VSD and PD conformations, channel-PIP2 interaction properties and VSD-PD coupling contacts, and (2) by applying Rosetta energy calculations to KCNQ1 VSD mutants to estimate the mutation-induced thermodynamic changes in protein stability, with results supporting the notion that protein destabilization is a common cause of mutation-induced KCNQ1 dysfunction. We anticipate that these Rosetta models will stimulate a new series of integrated structure-electrophysiology studies and personalized medicine approaches in KCNQ1 research.

## Results

### Model building and quality check of human KCNQ1 models

Structural models of human KCNQ1 (helix S0-S6) in the RC and AO state (**Fig. 1**) were generated by Rosetta homology modeling guided by the multiple sequence alignments shown in **Fig. S1** and **S2**. The template with the highest sequence identity (78%) to human KCNQ1 is the recently published cryo-EM structure of *X. leavis* KCNQ1 (PDB 5VMS) (27), which, however, appears to represent a decoupled state with an activated VSD and a closed pore (referred to as the AC state). Thus, the two domains served as separate templates for the AO and RC states, respectively. The *X. leavis* KCNQ1 VSD provided a template for the human KCNQ1 AO model whereas its PD was used in RC state modeling. An additional template for the open pore conformation in the AO model was derived from the crystal structure of the chimeric K_V_1.2-2.1 channel (PDB 2R9R) (31). Because no experimental resting state VSD structure of a voltage-gated potassium channel was available, templates of the VSD in the RC model were obtained from VSD structures of related non-mammalian proteins and another structural model: the resting VSD in *C. intestinalis* voltage-sensing phosphatase (Ci-VSP) (PDB 4G7Y) (28), VSD2 in the *A. thaliana* two pore calcium channel protein 1 (TPC1) (PDB 5DQQ) (29), and the resting VSD conformation C3 in a model of the Shaker channel (32).

**Figure 1:**
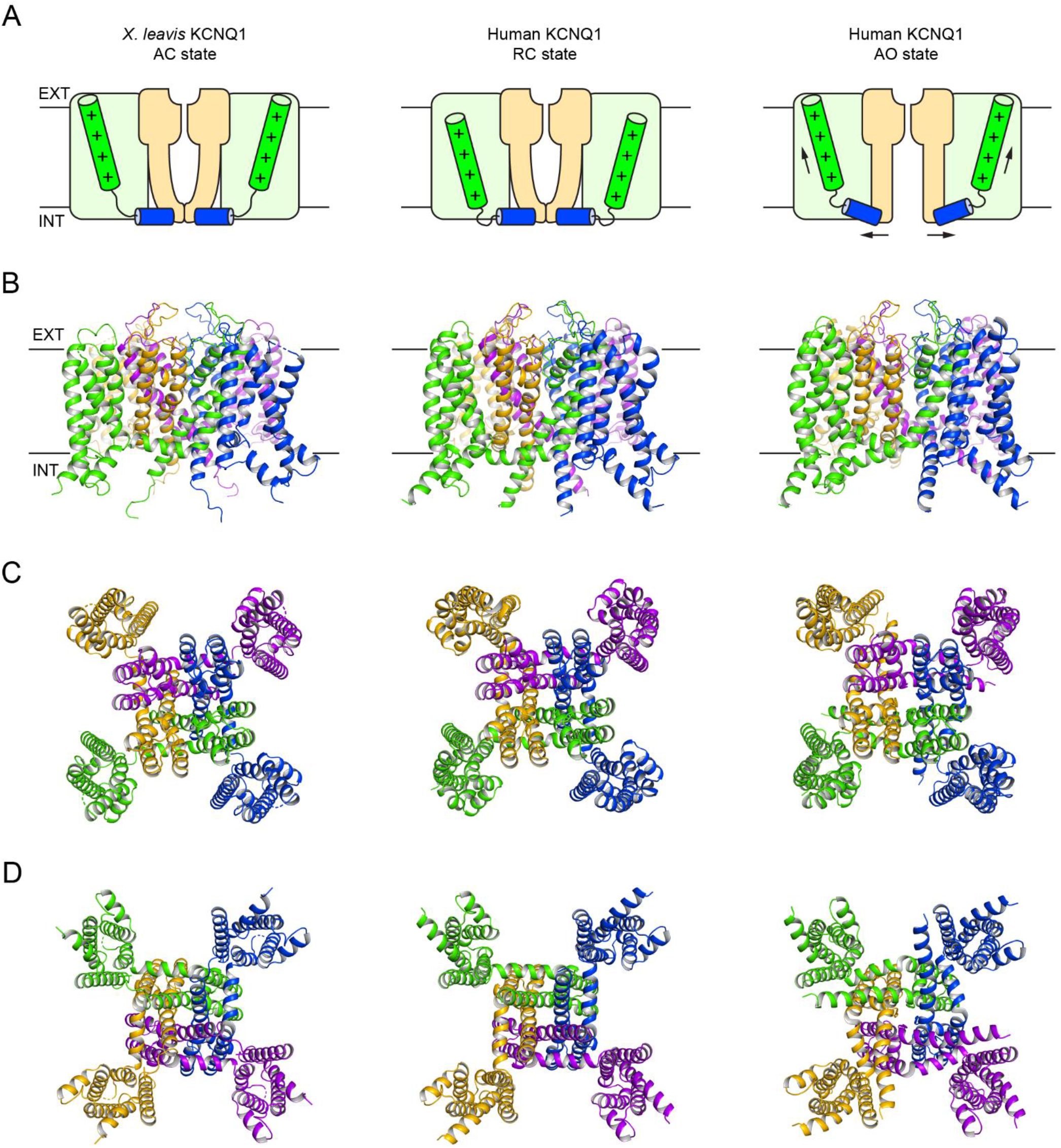
Overall architecture of the *X. leavis* KCNQ1 structure and human KCNQ1 homology models. **(A)** Schematic cartoon depicting the functional states of the VSD (light green box) and PD (light orange box). The S4 helix (with positive gating charges “+”) and S4-S5 linker helix are shown as green and blue cylinders, respectively. The *X. leavis* KCNQ1 structure is in a decoupled state with an activated VSD and closed PD. Homology models of human KCNQ1 were developed with the VSD / PD in the resting / closed (RC) and activated / open (AO) conformation. Black arrows illustrate conformational changes during voltage-dependent channel activation. **(B)** Side view of the KCNQ1 channel models. The approximate position of the membrane bilayer is indicated by vertical lines and the extracellular and intracellular sides are labeled EXT and INT, respectively. **(B)+(C)** View of the KCNQ1 channel models from the extracellular and intracellular side, respectively.

Rosetta homology modeling yielded an ensemble of low energy models that were assessed with Procheck (33) and Molprobity (34). The RC and AO models with the best Molprobity statistics (compare with **Table S1**) were selected as final models. They had favorable Molprobity and clash scores that were equal or better than the 90^th^ percentile of structures in the Molprobity database. No violations for backbone bonds or Cβ deviations were observed, and only four bond angles in both models fell outside of four times the standard deviation of dictionary values. In the AO model, all residues fell within favored (95.9%) or allowed (4.1%) regions of the Ramachandran plot. In the RC model, only 1% of the residues fell in disallowed regions with the remaining being in favored (92.8%) and allowed (6.2%) regions. Analysis of the side-chain conformations revealed 100% (AO model) and 99.5% (RC model) favored rotamers and no bad rotamers. Taken together these values are well within the range of high-resolution structural models and a positive indicator of the quality of our KCNQ1 models. PDB files of these models are provided in the Supporting Information (**S1 PDB** and **S2 PDB**).

### Structural validation of human KCNQ1 RC and AO models

Our KCNQ1 RC and AO models are displayed in **Fig. 1** and their topology is compared to that of *X. leavis* KCNQ1. KCNQ1 has a domain-swapped homotetrameric topology with the VSD of one subunit interacting with helix S5’ from a neighboring subunit. The VSD and PD in the AO model had a Cα-atom RMSD of 1.9 Å and 2.3 Å relative to the *X. leavis* KCNQ1 AC structure, respectively. For the RC model the Cα-RMSD for the VSD and PD were 4.1 Å and 1.0 Å, respectively. These values are considerably lower than those calculated for the first generation of human KCNQ1 models (16) which had RMSD values for the VSD and PD of 4.8 Å (3.5 Å when excluding the S2-S3 linker) / 3.0 Å (AO model) and 4.3 Å (3.6 Å) / 3.1 Å (RC model), respectively. Thus, the new KCNQ1 models are closer to the *X. leavis* KCNQ1 structure (i.e. the most closely related structural homolog available), which is a positive indicator for their improvement over the Rosetta models by Smith et al. (16) (see Discussion).

The KCNQ1 RC and AO models were validated rigorously against known experimental and structural data to assess model confidence. First, we calculated the channel pore radius and compared it to the closed PD conformation in *X. leavis* KCNQ1 (**Fig. 2**). In the RC model, the positions with the narrowest restriction along the ion conduction pathway are formed by residues G345, S349 and L353 which correspond to G335, S339 and L343 in *X. leavis* KCNQ1. The pore radius at those residues (~0.5 Å) is clearly below the ionic radius of a K^+^ ion (1.38 Å) confirming that the channel gate in the RC model is closed. In contrast, the channel gate is open in the AO model. The radius at the narrowest point of the pore at A344 is not smaller than 2.0 Å, which will allow a K^+^ ion to pass through. Beyond that point, the pore radius increases and remains larger than the K^+^ ionic radius. This transition point in the KCNQ1 pore coincides with the functionally important PAG motif, which corresponds to PXP in other K_V_ channels. The PAG motif serves as a hinge point allowing the S6 helix to bend and swing outward, which opens the channel gate. Mutations of the PAG motif abolish current but not channel surface expression (35) consistent with the idea that this region is important for movement of the activation gate. These observations for our Rosetta models were reproduced by pore radius measurements made for these models when employed in MD simulations (**Fig. S3**, see explanations below), further confirming the validity and stability of the PD structure in our KCNQ1 models.

**Figure 2:**
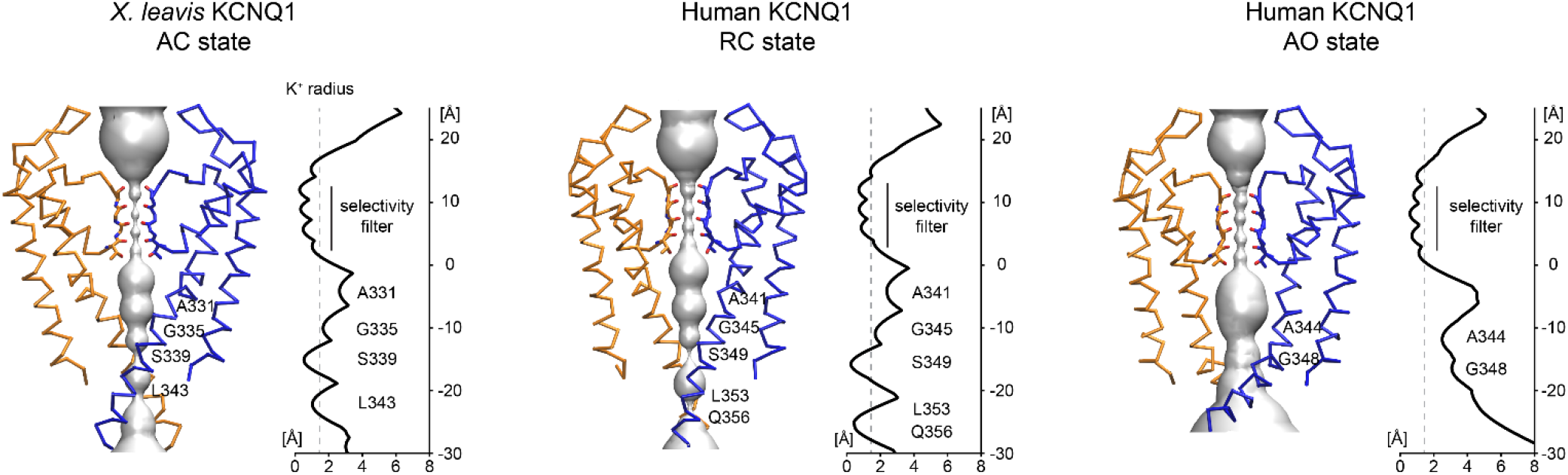
Pore conformation in the *X. leavis* KCNQ1 structure and human KCNQ1 RC and AO homology models. Left: surface representation of the channel pore. Subunits at the front and back are excluded for clarity. The selectivity filter is displayed as sticks. Amino acid residues facing the pore are labeled. Right: pore radius calculated from the *X. leavis* KCNQ1 structure and the human KCNQ1 homology models. The approximate radius of a K^+^ ion is indicated by a gray dashed line.

As a second validation, we measured state-dependent distances in our models between amino acid residues suggested to be responsible for voltage-dependent activation of the VSD and compared them to expectations from known residue pairings inferred from electrophysiological studies on charge-reversal VSD mutants of KCNQ1 (5, 6). Positively charged residues located along the S4 helix, commonly referred to as gating charges and labeled R1 through R6, undergo a series of stepwise transitions and successively pair with acidic residues on S2 and S3 that creates movement of S4 and confers voltage-sensitivity to the ion channel (36). Critical residues that coordinate S4 movement include E1 (E160/E150 in human/*X. leavis* KCNQ1), E2 (E170/E160) and D202 (D192 in *X. leavis* KCNQ1). In the resting VSD, E1 likely pairs with R1 (R228 in human KCNQ1) whereas the activated conformation is stabilized by an interaction between E1 and R4 (R237) (5, 6, 37). In addition, residue Q3 (Q234/Q224 in human/*X. leavis* KCNQ1) is found close to E2 in the cryo-EM structure of KCNQ1, which likely represents an activated state of the VSD. The described state-dependent residue pairings are correctly captured by our KCNQ1 Rosetta models and persistently detected in MD simulations of the RC and AO model (**Fig. 3**). The structural models also suggest additional interactions occur between helix S4 and residues of the charge transfer center: in the RC model R2 and Q3 interact with E2 and D202 whereas in the AO model H5 and R6 are close to those two acidic residues. Moreover, another source of structural stabilization of the VSD in its resting and activated conformation may come from interactions of the gating charges with membrane phospholipids. This idea is supported by the observation of frequent hydrogen bond contacts of R6 (RC) and R1 (AO) with phospholipids in the lower and upper bilayer leaflet during MD simulation (**Fig. 3**).

**Figure 3:**
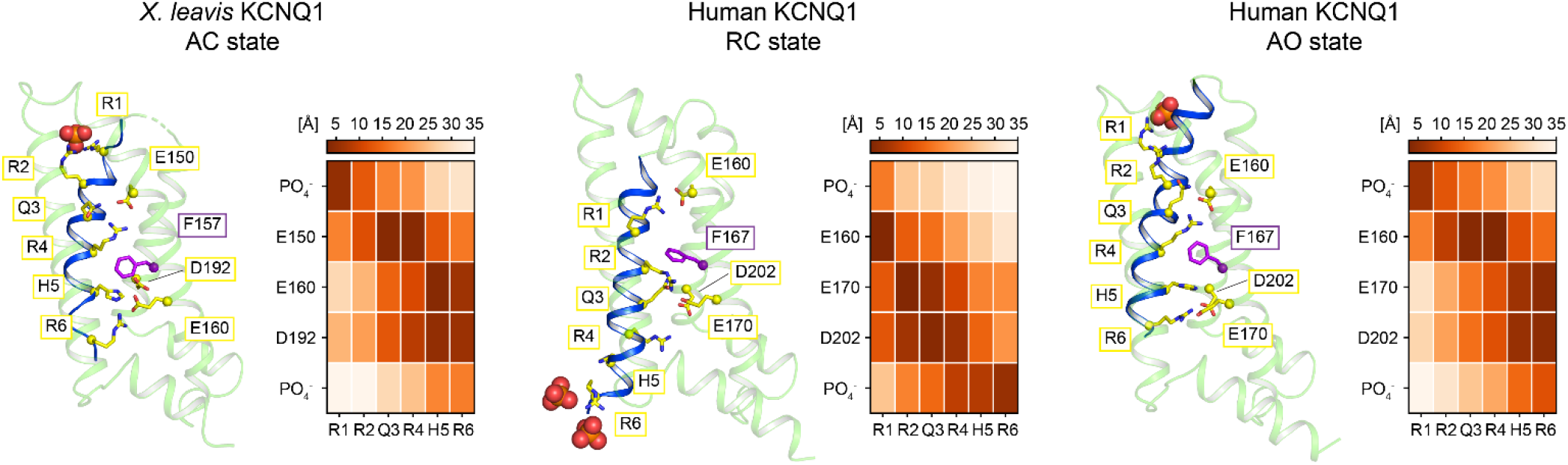
Voltage sensor conformation in the *X. leavis* KCNQ1 structure and human KCNQ1 RC and AO homology models. Left: VSD conformation; helices S0-S3 are colored pale green and helix S4 is colored blue. The side-chains of positively charged residues on S4 (yellow) and the gating charge transfer center residues E150/E160 (S2, yellow), F157/F167 (S2, magenta), E160/E170 (S2, yellow) and D192/D202 (S3, yellow) are depicted as sticks. Phosphate groups of phospholipids in the outer (top) and inner (bottom) bilayer leaflet are represented as spheres. Right: side-chain – side-chain distances between residues on S4 and gating charge transfer center residues. Values represent the average over four VSD copies, and were calculated from the cryo-EM structure of *X. leavis* KCNQ1 (after embedding in a POPC bilayer with CHARMM-GUI (95) and short minimization in Amber16 (41)) and from MD simulations of human KCNQ1 models. Distances were measured between the geometric centers of the side-chain atoms H_2_N=C_ζ_(NH_2_)-N_ε_H-C_δ_H_2_ (Arg), H_3_N_ζ_C_ε_H_2_ (Lys), C_γ_-N_δ1_-C_ε1_H-N_ε2_H-C_δ2_H (His), HOOC_γ_-C_β_H_2_ (Asp), HOOC_ε_-C_γ_H_2_ (Glu) and the lipid phosphate group PO_4_^-^.

Overall, the measured channel pore dimensions and the good agreement of residue pairings in the VSD with experimental data lend confidence to the reliability of our KCNQ1 structural models.

### Refinement of KCNQ1 models by MD simulation and study of KCNQ1-lipid interactions

Computational simulations have become an indispensable tool in the study of voltage-gated ion channels and have provided insights into different aspects of KCNQ1 function and regulation, such as KCNE1 binding (17, 38), drug (39) and lipid binding (40). Here, we assessed the structural stability of our KCNQ1 AO and RC homology models in all-atom MD simulations with Amber (41). We further validated that the simulations can reproduce known interactions of KCNQ1 with phosphatidyl-4,5-bisphosphate (PIP2) that we found to be in favorable comparison with experimental data. PIP2 is an important second messenger for cell signaling that binds to and regulates a wide variety of ion channels, including KCNQ1 (42). Depletion of PIP2 from the membrane suppresses VSD-PD coupling in KCNQ1 (23) but leaves the VSD activation intact.

A total of four RC state and four AO state simulations starting from the selected KCNQ1 models as well as three additional conformations from the ensemble of Rosetta homology models were conducted, each running for 400 ns. Simulations were performed under constant temperature and pressure conditions in explicit POPC bilayers that contained 10 mol% (~28 molecules) of PIP2 in the cytosolic leaflet (**Fig. 4A**). Overall, the KCNQ1 models remained stable in MD and deviated by not more than a Cα-RMSD of 4-5 Å from their respective starting conformations (**Fig. 4B** and **Fig. S4**). The RC and AO models maintained their closed and open pore structure (**Fig. 2** and **Fig. S3**), respectively, as well as their characteristic gating charge pairings (**Fig. 3**) as discussed above.

**Figure 4:**
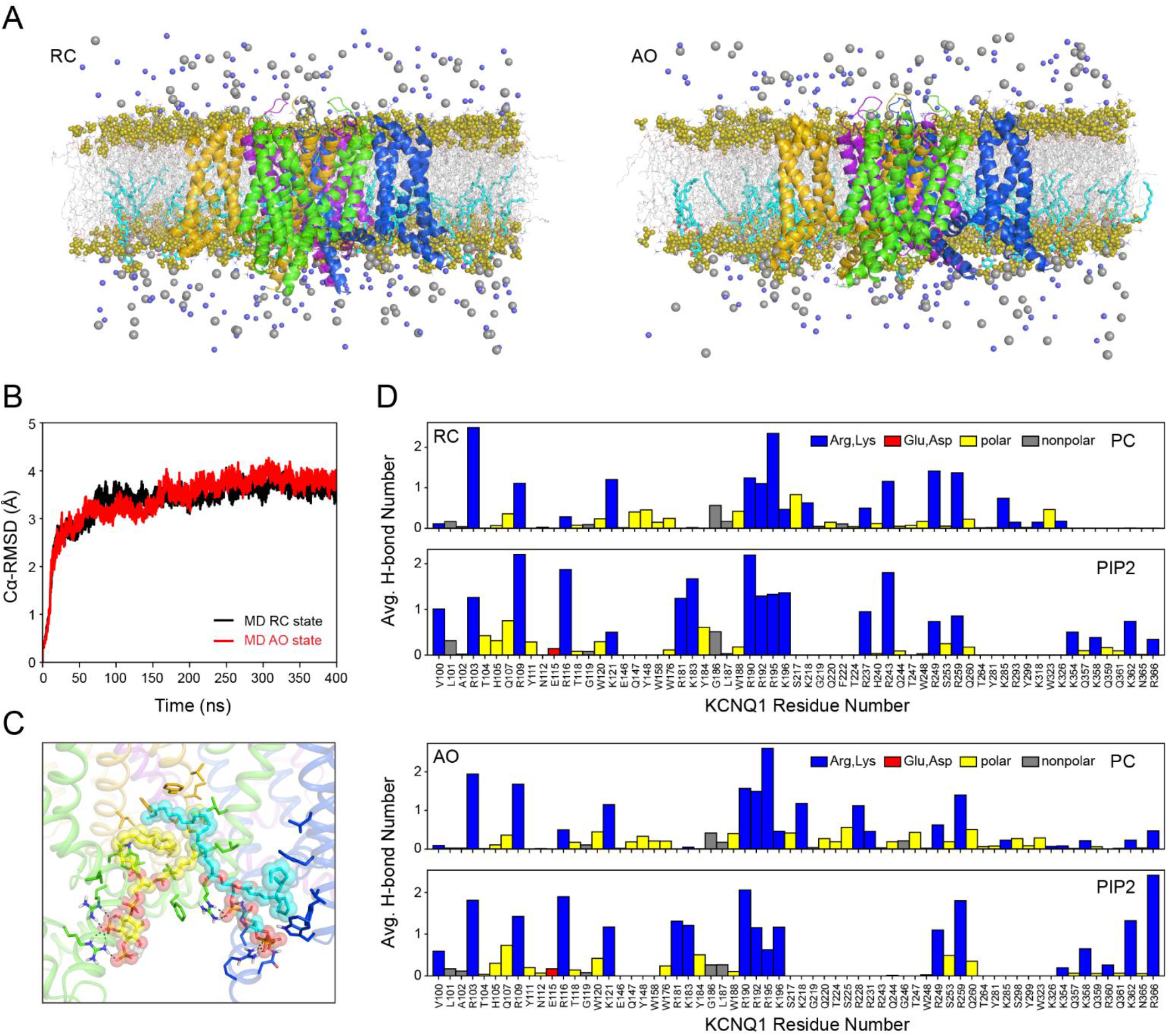
MD simulation of KCNQ1 homology models. **(A)** RC and AO models embedded in a POPC bilayer that contained 10 mol% PIP2 in the inner leaflet. The fatty acid chains of POPC and PIP2 molecules are depicted by gray and cyan sticks, respectively, and the head group phosphates are represented as olive spheres. Potassium and chloride ions are displayed as gray and light blue spheres and water molecules are excluded for clarity. **(B)** Cα-atom RMSD relative to the starting structure over the course of an MD trajectory for the RC and AO model. RMSD plots for all four RC and AO state MD simulations, respectively, are shown in **Fig. S4**. (C) Snapshot of the interaction of two PIP2 molecules with KCNQ1 as observed in MD. PIP2 molecules are drawn as spheres with carbon atoms colored yellow or cyan, respectively, and oxygen and phosphorous atoms colored red and orange, respectively. Different KCNQ1 subunits are colored green, blue, purple and orange. Side-chains of nearby KCNQ1 residues are depicted as sticks with heteroatoms and polar hydrogen atoms colored by their chemical identity (hydrogen: white, oxygen: red, nitrogen: blue). Potential hydrogen bond contacts are indicated by dashed lines. (D) Average number of hydrogen bonds per MD frame between KCNQ1 residues and phosphate groups in PC and PIP2 head groups calculated over the course of all RC and AO state MD trajectories.

The number of hydrogen bonds with POPC and PIP2 was analyzed after letting the system equilibrate for 100 ns. Because the mean square displacement values for lateral diffusion of POPC and PIP2 in Amber lipid simulations are in the order of several ten Å^2^ per 10 ns (43), this time span was long enough to allow the initially randomly placed PIP2 molecules to diffuse towards KCNQ1 and engage in close protein contacts (**Fig. 4C**). PIP2 was observed to bind preferentially in the cleft between two neighboring VSDs (**Fig. 4C**). It made numerous hydrogen bond interactions with residues on the S2-S3 linker (S2-S3L) (R181, K183, R190, R192), the S4-S5 linker (S4-S5L) (R249, R259) and the cytoplasmic end of S6 (S6_C_) (K354, R360, K362) (**Fig. 4D**) in agreement with the experimentally reported PIP2 binding sites (23, 24). Previous studies suggested that interactions of PIP2 with these mostly positively charged residues strengthen the coupling between the VSD and PD and are critical for pore opening (44). Consistent with its high functional importance, a series of LQTS-associated mutations affect the PIP2 binding region: K183R (45), K183M (46), R190Q (47), R190L (48), R190W (49), R192H (50), R192P (49), R259C (51), R259H (50), R259L (52), R360M (48), R360T (49).

Comparison of the locations of POPC and PIP2 interaction sites on the cytosolic juxtamembrane side of KCNQ1 (**Fig. 4D**) revealed some significant differences: POPC was unable to participate in hydrogen bonding with residues R181 and K183 in the short loop between S2 and the S2-S3 linker and with residues on S6c which are located within or below the water-interface region of the membrane. As a consequence of the short PC head group those residues are outside of the range of POPC but can still be reached by PIP2 that has a larger head group bearing two additional phosphates.

### High impact mutation sites map intra- and inter-subunit contact regions between the VSD and PD in KCNQ1 models

Using MD simulation, we further characterized the interactions between the VSD and PD in the RC and AO model (**Fig. 5**) and compared them to reported experimental data (26, 53–55). VSD-PD interactions translate the voltage-dependent movement of S4 mediated by leverage of the helical S4-S5 linker (S4-S5L) to channel opening and closing. Inference from studies of other K_V_ channels (56–58) and mutational scanning experiments on KCNQ1 (26, 53) have pointed to the interface between S4-S5L and the cytosolic end of S6 (S6_C_) as one important region for electromechanical coupling. In those studies, one group of mutations in S4-S5L (26) and S6_C_ (53) reduced the channel opening rate and shifted channel activation to more depolarized voltages whereas another group of mutations, specifically at V254 in S4-S5L and at L353 in S6_C_, prevented the channel from closing and led to a constitutively open channel. These results were taken to suggest that S4-S5L binds to S6_C_ to stabilize the closed state of the channel and that S4-S5L relocation in voltage-dependent activation releases tension on S6 that allows it to kink, promoting channel opening.

**Figure 5:**
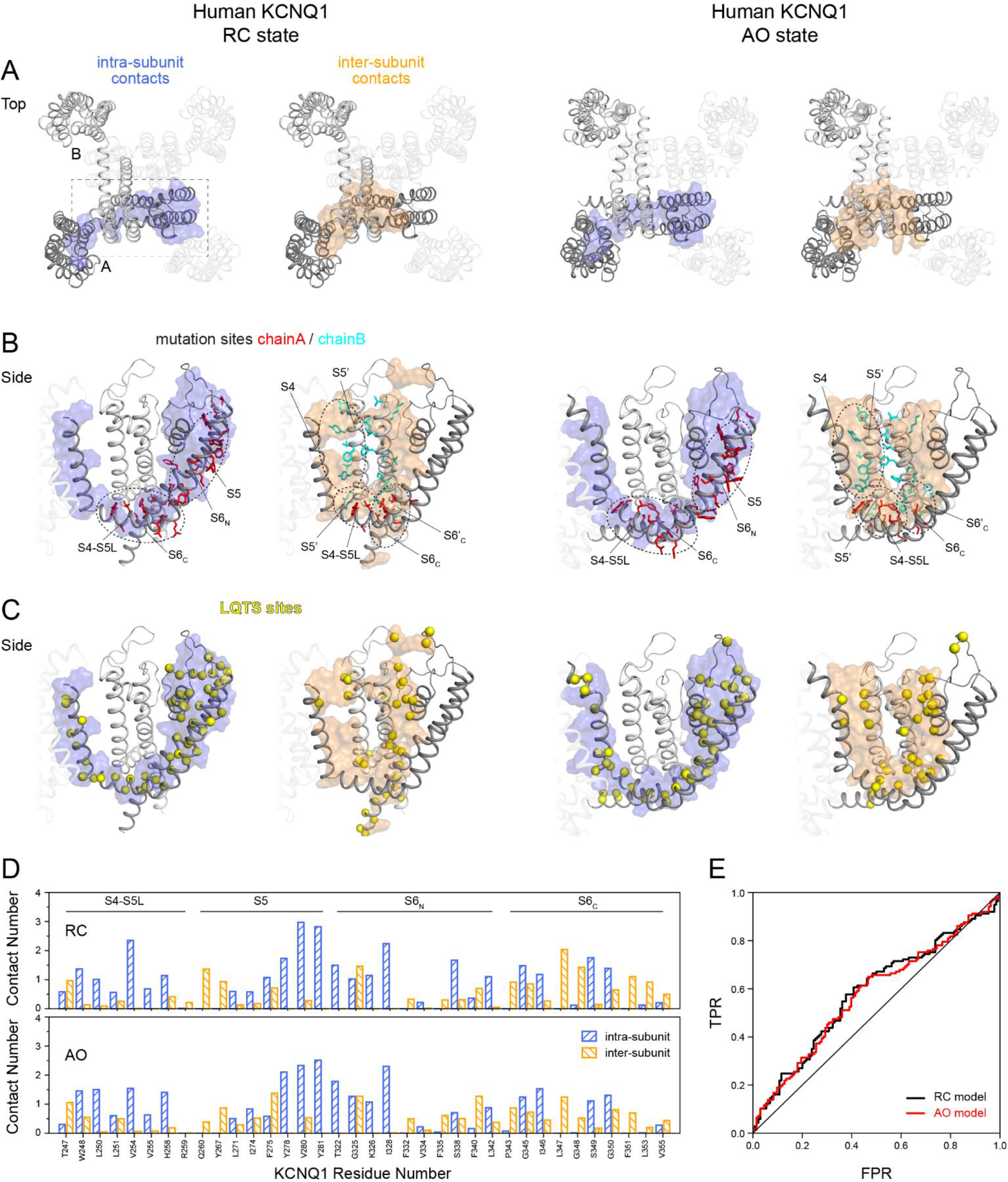
High impact mutations and LQTS sites fall within intra- and inter-subunit contact regions in KCNQ1 channel models. **(A)** Extracellular view of the KCNQ1 RC and AO models. For clarity, only two neighboring subunits are displayed while the other two chains are transparent. Residues in S4 to S6 that make contacts within the same subunit or with residues in the neighboring subunit are shown as surface representation and are colored light blue or orange, respectively. A cutoff value of 1.0 and 0.6 for the normalized contact number (i.e. mean total number of heteroatom contacts within a 4 Å distance divided by the number of heteroatoms of a given amino acid residue) of intra- and inter-subunit contacts was used for making the surface representations. The normalized contact number was calculated as the average over the four RC or AO state MD simulations, respectively (see Methods). **(B)** Residues corresponding to high impact mutation sites in KCNQ1 (26, 53, 55) located in S4-S5L, S5 or S6, respectively, are plotted onto the KCNQ1 models and depicted as sticks. Circled areas indicate intra- and inter-subunit contact regions. **(C)** Location of LQTS sites mapped on the KCNQ1 models depicted as yellow spheres. **(D)** Normalized contact number of residues at high impact mutation sites in S4-S5L, S5 and the N- (S6_N_) and C-terminal (S6_C_) end of S6. **(E)** Receiver operating characteristic (ROC) curve for classifying a KCNQ1 variant as LQTS or non-LQTS based on a residue’s contact number. The area under the curve (AUC) when using the RC and AO model was 59% and 58%, respectively.

In the RC and AO model, we identified extended regions of intra- and inter-subunit contacts (displayed in **Fig. 5A**) that tie the VSD and S4-S5L to the neighboring PD. Intra-subunit contacts were found between S4-S5L and S6_C_ and between S5 and the N-terminal end of S6 (S6_N_). Inter-subunit contacts were observed for S4 and S4-S5L with S5’ from an adjacent KCNQ1 chain and between two S6 helices from neighboring chains that cross each other near the channel gate. Strikingly, mutation sites in S4-S5L, S5 and S6 with high impact on channel opening and closing (26, 53, 55) are clustered in those contact regions and have consistently high contact numbers (**Fig. 5B** and **5D**) indicating that those regions are important for channel function. For clarity, mutation sites in the selectivity filter and P helix were not marked in **Fig. 5B**. Those elements share only a small contact area with the VSD and many mutations cause complete channel loss of function (LOF) because of the detrimental effect that these mutations have on the structural integrity of the selectivity filter and extracellular pore entrance gate.

In the RC model, the S4-S5L – S6_C_ contact surface is formed by residues L250, L251, V254, V255, H258 on S4-S5L and I346, L347, G350, F351, L353 and K354 on S6_C_. The side-chains of V254 and L353 were found to be proximal to each other in the model consistent with the results of combinations of mutations at those two sites (26) that were able to rescue channel function. Interestingly, L353 also engages in side-chain – backbone interactions with one of its symmetry mates in the neighboring subunit. As viewed from the cytosolic side, the four symmetric copies of L353 are arranged circularly with the orientation of their side-chains directed clockwise. L353 is located one helical turn below the helix-helix crossing point at S349, which forms the narrowest constriction of the channel pore (compare with **Fig. 2**). Hence, our structural model can provide a conclusive explanation for the constitutively open channel phenotype of KCNQ1 mutants with charged amino acid substitutions (K/E) at L353 (53): steric repulsion between close local charges at the position of L353 pushes the S6 helices apart and prevents the channel from closing.

The contact region between S4-S5L and S6_C_ is maintained in the AO model as well and comprises the same group of residues but is reoriented slightly when S4-S5L and S6_C_ slide against each other during pore opening. The L353 sites, however, become separated from each other when S6 is kinked and are not in contact in the AO model.

The inter-subunit contacts span similar regions in the RC and AO model, respectively, but differ in their local structure reflecting the conformational changes that occur during channel activation as described below. Inter-subunit contacts are probably also the reason for the global motions in KCNQ1 that we observed as characteristic regular patterns in the dynamic residue cross-correlation matrix (DCCM) (**Fig. S5A**), and by performing principal component analysis (PCA) on the intra-membrane segments in KCNQ1 (**Fig. S5B+C**). The global motions of KCNQ1 in MD simulation feature a rigid-body-like swing movement of the VSDs that includes both an ‘in-membrane-plane’ as well as an ‘out-of-membrane-plane’ component. Two neighboring VSDs were observed moving antiparallel to each other whereas the two domains on opposite sides of the tetrameric channel moved in the same direction (see **Fig. S5** for further details).

We further noticed that many mutation sites that have been associated with LQTS fall within intra- and inter-subunit contact regions in KCNQ1 (see **Fig. 5C**). A simple classifier that distinguishes LQTS from non-LQTS sites based on residue contact number yields a receiver operating characteristic (ROC) with an area under the curve (AUC) of 59% and 58% for the RC and AO model, respectively (**Fig. 5E**). This shows that LQTS sites are enriched in areas with high contact number and that mutations at those sites are critical to ensure channel function.

### Transition from the RC to the AO state triggers changes in inter-subunit contacts between the VSD and pore

Comparison of the RC and AO model revealed conformational changes at the VSD-PD interface and in the PD (**Fig. 6**) (in addition to S4 in the VSD), which likely reflect the changes happening during depolarization-activated channel gating. Triggered by the gating charge movement in the VSD, S4 slides along S5’ and moves upwards by about two helical turns. Hydrophobic residues on the S5’-facing side of S4 swap their positions and come into contact with new residues on S5’ while S4 is moving (compare with left plot in **Fig. 6A**). The S4-S5L is pulled upwards as well and engages in new interactions with residues located two to three turns higher on S5’. For instance, in the RC model, T247 makes contacts with the first residues in S5’ (Q260, E261) but is packed against T264, Y267 and I268 in the AO model. W248 makes few interactions with S5’ in the RC model but makes many putative stabilizing contacts with Y267, I268 and L271 in the AO model (**Fig. 6B**). Moreover, movement of S4-S5L allows S6 to kink and move sideward cantilever-like. This movement breaks contacts between the ends of S6_C_ and induces a crosswise sliding of the S6 helices (compare with right plot in **Fig. 6A**) that widens the channel pore diameter. Specifically, residue F351 slides upwards by about two helical turns and becomes tightly packed in the AO model where it appears to bridge sites on S4-S5L with S6_C_ of the same and S6’_C_ of the neighboring subunit (**Fig. 6B**). The observation that F351 is at a critical spot in KCNQ1 with many contacts to other residues is consistent with the fact that F351 is highly conserved in K_V_ channels, and that it is a high impact mutation site (53). Experimental studies (53) showed that the F351A mutant featured a rightward shifted current-voltage relationship and longer activation and deactivation times that mimicked the I_KS_-phenotype of KCNQ1, and was ascribed to a destabilization of the KCNQ1 intermediate/open state (6) similar to the effects of KCNE1.

**Figure 6:**
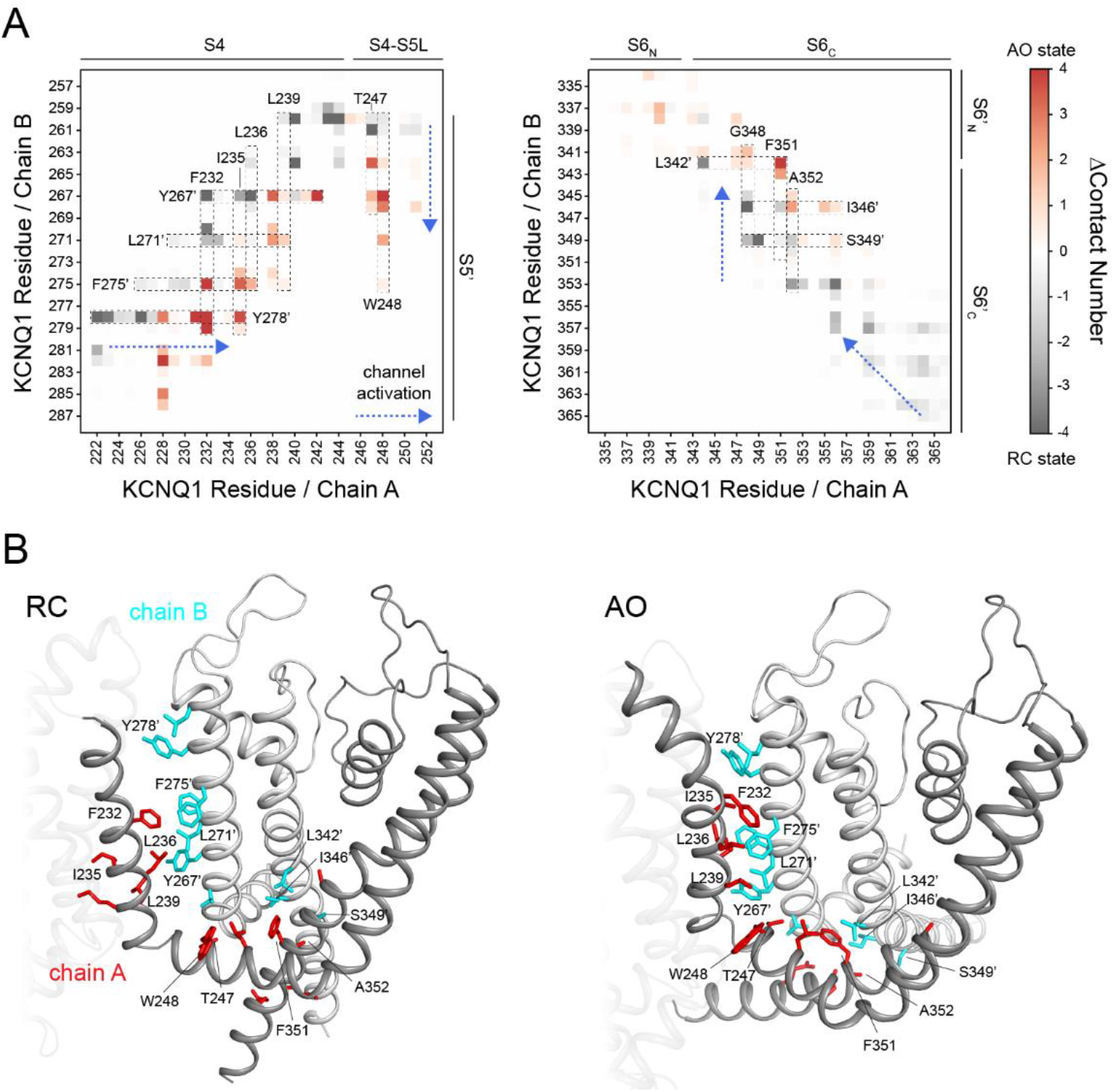
Changes of inter-subunit contacts in KCNQ1 during channel activation. **(A)** Average number of heteroatom contacts between residues in two neighboring KCNQ1 chains A and B. The contact number was defined as number of heteroatom pairs within a 4 Å distance and averaged over the last 300 ns of MD and all four pairs of neighboring subunits in the KCNQ1 tetramer. The section of the contact matrix corresponding to the interface of S4 and S4-S5L with S5’ (left) and of two neighboring helices S6 and S6’ (right) is shown. A gray color means this contact is observed in the RC model whereas a red color denotes a contact formed in the AO model. Changes in specific residue contacts occurring while the channel transitions from the RC to the AO state are framed and labeled by their corresponding amino acid residue. The direction of the structural changes with channel activation is indicated by a blue arrow. **(B)** Cartoon representation of the inter-subunit interface in the RC (left) and AO (right) model, respectively. For clarity, only helices S4 to S6 are shown. Residues which are part of the inter-subunit interface and fall within regions of the contact matrix in **(A)** are depicted as sticks. Residues with drastic changes in their contact pattern as identified in **(A)** are labeled.

In conclusion, we found that many high impact mutation sites map to interactions between the VSD and PD that undergo state-dependent changes when the channel transitions from the RC to the AO state illuminating the importance of those interactions in electromechanical coupling.

### Rosetta-predicted stability changes in KCNQ1 models correlate with experimental channel loss of function data

While progress in the functional characterization of LQTS-associated mutations has been made (12–14) the mechanistic molecular basis of channel dysfunction for most KCNQ1 mutations is still unclear. The dearth of direct experimental data that can help to determine how mutations alter KCNQ1 structure and function has prompted computational modeling approaches (16, 59).

Here, we assessed the capability of our structural models to predict the change in thermodynamic stability (ΔΔG) in KCNQ1 variants and whether the predicted energy changes can account for alterations in channel function. We used a Rosetta ΔΔG prediction protocol (60) allowing for flexibility in the protein backbone and side-chains around the mutation site. Protein backbone flexibility was modeled by a series of backrub moves (61) that rotate short main-chain segments (three to twelve residues) as rigid body about an axis defined by the starting and ending Cα-atom of the segment. This aims at alleviating the prediction for mutations with a large change in amino acid size. Side-chain flexibility was modeled using discrete rotamer conformations and simulated annealing (i.e. side chain packing). The ΔΔG was calculated as mean difference between the three top-scoring models of wild type (WT) and mutant KCNQ1 and averaged over ten different starting conformations taken from the ensemble of Rosetta homology models as described in Methods.

ΔΔG predictions were made for a set of single-site KCNQ1 variants with the affected sites located in the VSD (shown in **Fig. 7A**, and divided by functional classes in **Fig. S6**). These KCNQ1 VSD variants were previously biochemically, biophysically, and functionally characterized (15), which provided the mechanistic basis for their pathogenicity. Not included from this original set of 51 KCNQ1 variants was the deletion mutant ΔF167 because modeling of deletions and insertions is currently not supported by the Rosetta ΔΔG prediction protocol. Furthermore, ΔΔG predictions for mutations to proline were found to be off-scale due to incompatible backbone torsions yielding Ramachandran and ring closure penalties as noted earlier (62). Therefore, their ΔΔG values were considered less reliable and also not included in the analysis leaving ΔΔG data for 44 single-site KCNQ1 variants.

**Figure 7:**
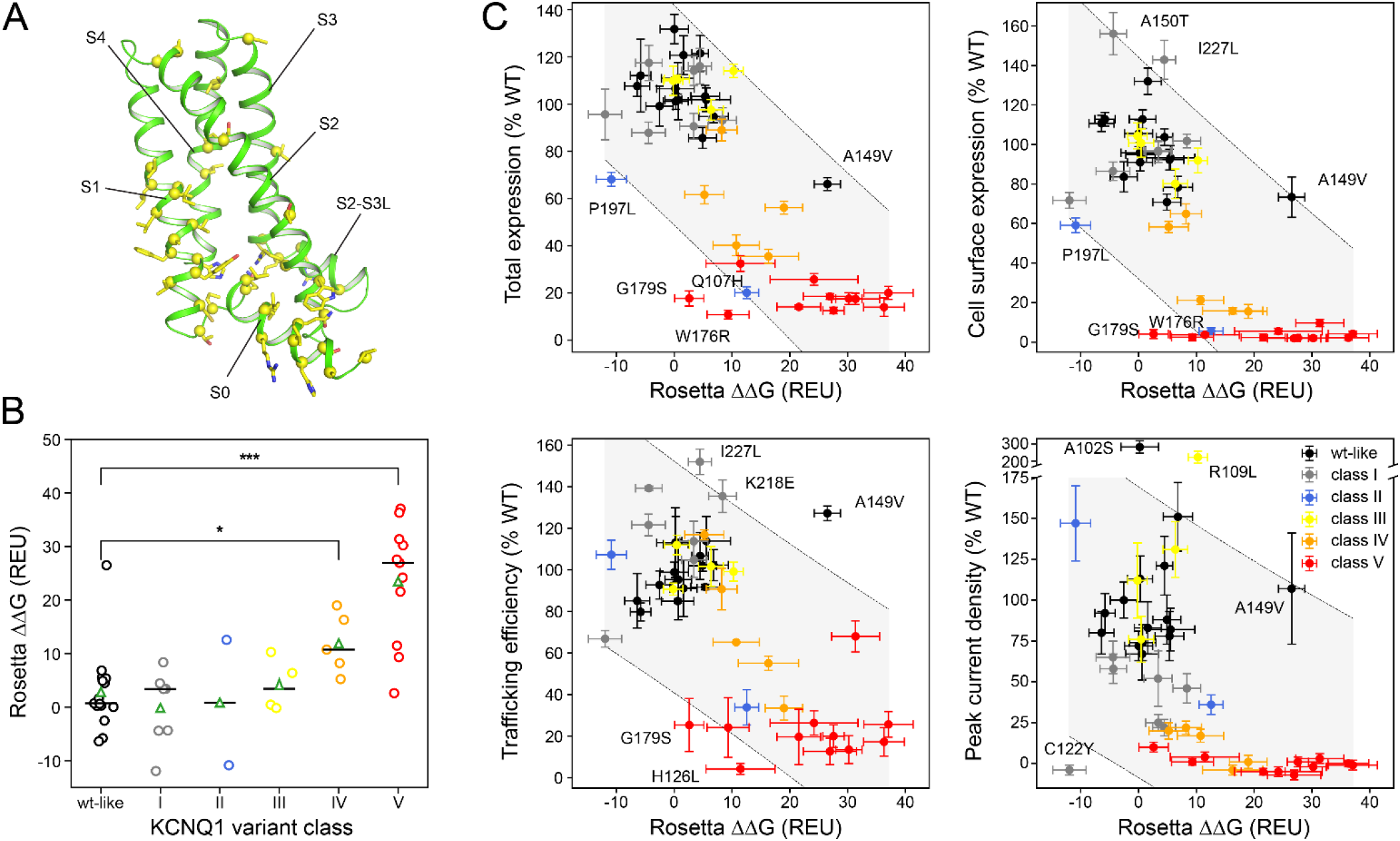
Computationally predicted stability changes of KCNQ1 variants with mutations in the VSD. **(A)** Location of mutation sites in the KCNQ1 VSD. Backbone sites are mapped by yellow spheres and the native amino acid residue is indicated by yellow sticks. **(B)** Distribution of Rosetta ΔΔG values for the six functionally distinct classes of KCNQ1 VSD variants (15) (i.e. WT-like and classes I to V for non-functional variants) calculated with the AO homology model. The median and average values are drawn as a black horizontal line and a green triangle, respectively. The median of class IV and V is compared to that one of WT-like variants using a Kruskal-Wallis H-test (* p < 0.05, ** p < 0.01, *** p < 0.001, n_WT-like_ = 15, n_IV_ = 5, n_V_ = 11). ΔΔG values for mutations to proline are off-scale (ΔΔG_L114P_ = 70.5 ± 4.1 REU, ΔΔG_L131P_ = 76.0 ± 3.0 REU, ΔΔG_L134P_ = 68.3 ± 4.1 REU, ΔΔG_R195P_ = 56.8 ± 2.0 REU, ΔΔG_Q234P_ = 71.0 ± 5.0 REU, ΔΔG_L236P_ = 65.8 ± 6.0 REU) due to incompatible backbone torsions in the starting model yielding bad backbone and proline ring geometries and were not used in the analysis. **(C)** Correlation plots of total expression level, cell surface expression, trafficking efficiency and channel peak current density versus calculated Rosetta ΔΔG values (mean ± S.E.M.). KCNQ1 variant classes are indicated with different colors. Variants falling outside of or being close to the boundary of the 95% confidence interval for a linear regression model (gray shaded area) are labeled and their structural models are shown in **Fig. S9** in the supplement. The experimental data and mutant classification scheme are from reference (15) and are listed together with the computed ΔΔG values for RC and AO models in **Tab. S2**. Class I variants traffic normally but exhibit channel LOF. Class II mutations induce mistrafficking, but do not alter the functionality of the small population of correctly-trafficked channel. Class III variants traffic normally and are conductive, but exhibit altered channel regulation. Class IV mutations cause mistrafficking and channel LOF. Class V variants are expression and trafficking-defective and have so low current that channel function is unmeasurable.

**Figure 7B** shows the distribution of ΔΔG values for those variants calculated with the KCNQ1 AO model as a function of variant class. Variants were previously assigned one of six classes based on their channel phenotype: WT-like and classes I-V for variants that have perturbations in the expression level, surface trafficking, and/or channel current density (see also the caption for **Fig. 7** and the footnote of **Tab. S2** for a description of the class phenotype). Roughly, the severity of the channel LOF increases from class I to V with variants in class IV and V having the most severe perturbations. The median ΔΔG value of these two classes was significantly higher (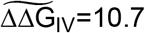 REU (p_IV_ < 0.05), 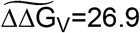 REU (p_V_ < 0.001)) than that one of WT-like variants 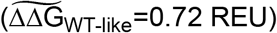 indicating the mutant protein is thermodynamically destabilized. It must be mentioned that Rosetta ΔΔG values (in Rosetta Energy Units, REU) do not necessarily have the same scale as experimentally measured protein stability values (in kcal/mol), but that their relative order (e.g. between different variant classes) is more instructive.

The predicted ΔΔG values were correlated with the expression, trafficking and current density data (**Fig. 7C**) suggesting protein stability is an important factor that contributes to these channel properties, and protein destabilization is a main cause of channel LOF. This idea is supported by NMR measurements on the KCNQ1 voltage sensor (15), which for two of seven class IV variants and ten of 14 class V variants had revealed spectra that were characteristic for a folding-destabilized protein. Another four class V variants failed to express the voltage sensor in *E. coli* also indicating protein folding defects (15). The channel state of the structural model (RC or AO) had no appreciable influence on the ΔΔG predictions. Similar results were also obtained with the RC model (**Fig. S7** and **S8**), indicating that the investigated mutations affected both channel states to a similar degree.

The correlation plots in **Fig. 7C** seem to have a complex, non-linear dependency, specifically the plot for peak current density, suggesting additional factors other than protein stability affect the named properties and/or that some variants had incorrect ΔΔG predictions. Variants that fell outside the 95% confidence interval for a linear model included four class I variants (C122Y, A150T, K218E, I227L), three class V variants (G179S, W176R, H126L), A102S and A149V (both WT-like) as well as P197L (class II) and R109L (class III). Structural models for these variants are shown in **Fig. S9** in the supporting information and their experimental and ΔΔG data are listed in **Tab. S2**. In particular, it was noticeable that class I variants had ΔΔG values (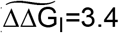 REU or 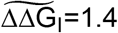 REU with the AO or RC model, respectively) not significantly different from WT-like variants 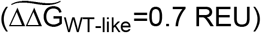, which failed to explain why these channels have low current density. For example, C122Y was predicted a negative ΔΔG (i.e. more stabilized than WT) but this channel was nonconductive. Two other variants for which channel function was not correlated with thermodynamic stability were A102S and R109L which had normal (0.2 REU) or increased ΔΔG values (10.3 REU), respectively, but exhibited current densities more than two times larger than WT (i.e. these were gain-of-function mutations).

The other few cases where the stability prediction was inconsistent with the functional data were likely due to incorrect ΔΔG calculations, which was supported by visual inspection of the respective models (see **Fig. S9**). For instance, A149V was predicted a high positive ΔΔG but had only moderately reduced expression levels and a trafficking efficiency higher than WT. Visual inspection of the structural model of this variant (**Fig. S9**) revealed remaining steric clashes between the valine side-chain and the nearby protein backbone causing a high repulsive score and indicating insufficient backbone conformational flexibility in modeling. G179S, W176R and H126L had ΔΔG values that were lower than other class V variants, mostly due to score balancing interactions made by the substituted amino acid residue in the mutant model, inconsistent with their low expression levels, trafficking and conductance.

## Discussion

### Improvements of upgraded KCNQ1 Rosetta models

Our ability to improve homology models of human KCNQ1 was largely informed by the new *X. leavis* KCNQ1 template, the closest structural homolog available. Consequently, the Rosetta models presented here bear high similarity to the *X. leavis* KCNQ1 cryo-EM structure and are closer to it than the Rosetta models by Smith et al. (16). The backbone RMSD for the VSD / PD compared to *X. leavis* KCNQ1 is 1.9 Å / 2.3 Å (this study) and 4.8 Å / 2.9 Å (Smith et al. (16)), respectively. For the closed state, the RMSD values for the VSD / PD are 4.1 Å / 1.0 Å (this study) and 4.3 Å / 3.1 Å (Smith et al. (16)), respectively. This indicates considerable remodeling changes were applied to the Rosetta models: comparison of the models presented herein and in Smith el al. shows an RMSD for the VSD / PD of 5.3 Å / 3.2 Å (closed) and 5.2 Å / 2.4 Å (open). It is important to mention that the Rosetta models of this work also compare favorably with their other structural templates: the PD in the AO model with the PD in K_V_1.2-2.1 (1.9 Å) and the VSD in the RC model with the VSD in Ci-VSP (3.6 Å for sequence-aligned regions; 2.6 Å for S1+S2) and TPC1 (4.4 Å for sequence-aligned regions; 2.1 Å for S1+S2). These results, together with the favorable stereochemical properties of the new human KCNQ1 Rosetta models and their valid VSD and PD conformations as illustrated in Results, lends confidence that they represent a substantial improvement over the preceding models. It is also important to point out that the Rosetta models by Smith et al. did not include the S0 helix in the VSD, which is now appreciated to represent a crucial element for structural stability (see reference (15) and Results on KCNQ1 VSD mutations). The previous models further fail to satisfy the E1-R1 (E160-R228) gating charge interaction in the RC state which is an experimentally established contact stabilizing the resting state (5, 6), however, was not known for KCNQ1 at the time of the first model generation.

The improved Rosetta models should be regarded as working models that are subject to revision as more experimental structural data become available. Specifically, we need to point out a few limitations. Of lower confidence are those regions for which no coordinate information in the template structures was available, requiring *de novo* modeling. Furthermore, higher uncertainty applies to transition regions where templates from different sources were hybridized. Lower confidence regions include the loop between S4 and S4-S5L in the RC and AO models, the loop between S3 and S4 in the AO model and the loops between S0 and S1 and between S2-S3L and S3 in the RC model.

### Implications for KCNQ1 structural biology questions

Given improved completeness and accuracy, the Rosetta models provide helpful tools to refine and advance our understanding of the structure-function relationships in KCNQ1. A hallmark of the KCNQ1 channel is its distinctive functional properties observed upon association with different members of the family of KCNE accessory proteins. While KCNE1 increases current density and delays channel opening and closing (8, 9), KCNE2 constitutively activates KCNQ1 and reduces current levels to approximately 50% of KCNQ1 alone (63). KCNE3 also makes KCNQ1 constitutively open but increases current by ~10 fold (22, 64).

The mechanism by which KCNE1 modulates KCNQ1 is a longstanding yet still unsettled debate and different models have been put forward such as alteration of the VSD movement (65), perturbation of pore opening (66) and changes in VSD-PD coupling interactions (6). A mapping onto our KCNQ1 Rosetta models of putative KCNE1 binding sites identified in cysteine crosslinking (18, 19, 66–68), mutational scanning and double mutant cycle (21, 69) experiments (see **Fig. S10**) suggests very similar binding pockets for KCNE1 in the RC and AO state. Preliminary docking calculations with our Rosetta models also suggest that KCNE1 makes similar interactions with KCNQ1 in the RC and AO states and adopts similar orientations, which is an unexpected result and was not observed in the original Rosetta models (17) indicating that a revision of the old KCNQ1/KCNE1 models is well-merited.

Of the many previous structure-function studies of KCNE1, we want to highlight here the model proposed by Zaydman et al. (6) who suggested that KCNE1 modulates KCNQ1 function by changing the VSD-PD coupling interactions. This idea stems from observations of KCNE1 inhibiting channel opening when the VSD is in the intermediate conformation between resting and activated state, as well as studies of F351A, a mutation that impairs coupling by weakening the contact between S4-S5L and S6_C_, mimicking the KCNE1-phenotype of KCNQ1. Combining electrophysiology and kinetic modeling, Zaydman et al. concluded that changes in state-dependent VSD-PD interactions may explain many of the effects of KCNE1 including its delay of channel opening, increased channel conductance and pharmacological properties. Consistent with this idea, we identified extended intra- and inter-subunit contact regions between the VSD and PD in our Rosetta models (**Fig. 5**) that potentially contribute to electromechanical coupling in addition to the established S4-S5L – S6_C_ contact. Channel activation features state-dependent changes in those VSD-PD interactions, specifically in inter-subunit contact regions (**Fig. 6**). We speculate that KCNE1 modifies VSD-PD interactions, changing the properties of the open pore conformation and affecting the channel open probability. Previous docking studies (17, 38, 69) converged on a model in which KCNE1 is bound in a crevice between the VSD and PD of two neighboring subunits. The intimate interactions that KCNE1 makes in these models with several sites on S4, S4-S5L, S5 and S6 seems well suited to modulate electromechanical coupling interactions.

Explicit structure-function knowledge concerning KCNE2 binding and regulation of KCNQ1 is limited. Results from cysteine scanning (70), double mutant cycle analysis and computational docking (21) suggested a different interaction mode for KCNE2 than for KCNE1. However, a widely recognized molecular model for the KCNQ1-KCNE2 interaction has yet to be established. The KCNQ1 Rosetta models provide a means to devise new experimentally-testable hypotheses on KCNE2’s mode of action that should trigger further progress in this field.

For KCNE3, a series of functional studies (71, 72) have converged on the appreciation that it acts primarily by stabilizing the VSD, especially S4, in its fully activated state, thereby increasing the open probability at resting potential and making KCNQ1 a constitutively open channel. The stabilization of S4 arises from salt bridge interactions between R228 on S4 and residues D54 and D55 on KCNE3 (71) and possibly by another polar contact between Q244 on KCNQ1 and R83 on KCNE3 (22). Our Rosetta model of the AO state supports this mechanistic explanation by showing R228 and Q244 at surface-accessible sites at the extracellular and cytosolic ends of S4 where those residues extend into the inter-subunit crevice formed between the VSD and PD. Binding of KCNE3 to R228 on the extracellular side and to Q244 on the cytosolic side is suggested to lock the S4 helix in place and stabilize the activated VSD conformation. Recent studies also suggest an additional effect of KCNE3 on the PD (73) which can be investigated with the new KCNQ1 models e.g. through docking and MD simulations.

### Potential use of KCNQ1 models for personalized medicine

Substantial evidence suggests a majority of disease-causing mutations act, at least, by destabilizing the folded conformation of the encoded protein (74, 75) thereby reducing the population of the functional form of the affected protein in cells. Predicting the phenotype of a mutation and interpreting variants of unknown significance discovered by genome sequencing data can greatly benefit from methods to quantitatively compare the stabilities of WT and mutant forms of proteins. For water-soluble proteins, computational algorithms are able to predict mutation-induced stability changes with reasonable accuracy (76). However, these methods perform significantly worse when applied to membrane proteins (77), highlighting the pressing need to develop methods with improved performance for membrane proteins.

In light of personalized treatment strategies for LQTS, the KCNQ1 Rosetta models will be broadly useful for interpreting the functional phenotype of KCNQ1 variants that we demonstrated here by predicting the energetic change in protein stability upon mutation compared to WT KCNQ1. Of the 44 single-site missense mutations studied here, 25 were LOF mutations (channel current <65%WT). Huang et al. (15) found 18 of these LOF mutations exhibited lower (<65%WT) cell surface expression than WT and 13 were severely folding-destabilized as indicated by their NMR spectrum or the inability of the VSD to express in *E. coli*. For these subsets of variants, we found the median ΔΔG was 17.7 REU and 24.2 REU, respectively, which translates to 13 of 18 and 11 of 13 variants having largely increased ΔΔG values above 10 REU (**Fig. 7C**). In contrast, the median ΔΔG of the 19 non-LOF variants was significantly lower (0.7 REU, p<0.001), and for the 26 variants with WT-like cell surface expression the ΔΔG was 1.1 REU (p<0.001). Thus, the Rosetta ΔΔG predictions support the conclusion that the KCNQ1 LOF variants were energetically destabilized and that protein destabilization represents the likely cause for protein misfolding and low cell surface expression. We previously noticed (15) many class IV and V variants, which are folding-destabilized, have mutation sites located in S0 and S0-contacting regions (see **Fig. S6**). The Rosetta models and MD simulations revealed a network of interactions of residues in S0 with residues in S2 and S4 that appear to glue the VSD helix bundle together. This observation illuminates the role of S0 as an important scaffolding element in KCNQ1 and potentially also in other K_V_ channels.

Energetic destabilization and trafficking defects is a common disease mechanism associated with gene mutations that impact membrane proteins (78). Examples include the HERG potassium channel and cardiac arrhythmia (79), the human peripheral membrane myelin protein 22 (PMP22) and Charcot-Marie Tooth disease (80), and a subset of mutations in the cystic fibrosis transmembrane conductance regulator (CFTR) protein (81). It is likely that mutation-induced destabilization is also a trigger for the disease-causative intracellular retention and/or degradation of other membrane proteins e.g. myelin proteolipid (82).

Noticeable exceptions of variants for which Rosetta-predicted changes in protein thermodynamic stability were not correlated with KCNQ1 loss- or gain-of-function were especially variants in class I (e.g. C122Y) as well as A102S (WT-like) and R109L (class III). C122Y is a LOF variant and associated with LQTS(83). For this case, the reason for LOF is unknown. In our KCNQ1 models, C122 is close to the putative KCNE1 binding site in KCNQ1 between S1 and S5’ of two neighboring subunits. One may speculate mutation-induced changes of the local structure around C122 could affect the presentation of side-chains on S1 towards KCNE1 and therefore affect KCNE1 binding. Alternatively, the C122Y mutation could affect S4 movement through an allosteric mechanism. Five other mutations in class I (with normal or higher expression and trafficking levels but low or no conductance) are located in regions that are critical for KCNE1 binding and channel gating: on S1 (L134P), S4 (I227L, Q234P) and on S4-contacting regions (A150, K218). L134 is lipid-exposed and its side-chain is pointed towards the putative KCNE1 binding site. I227 is one position before the first gating charge R1 while Q234 occupies the third gating charge position (R3 in other channels, Q3 in KCNQ1). A150 is located in the S1-S2 loop and K218 in the S3-S4 loop, both of which undergo varying interactions with S4 during VSD activation. Thus, the unusual behavior of these class I variants may be related to the location-specific role these residues play in voltage-dependent channel activation mechanism.

The reason for increased conductance of A102S and R109L is unknown. Both sites are located in the first half of S0 and are lipid exposed. A structural mechanistic explanation for them is currently hampered by the lack of structural data for the N-terminal domain before S0, which is not part of the Rosetta models.

Knowledge of KCNQ1 LOF mechanisms pertaining to a given patient may soon facilitate the design of personalized treatment strategies for LQTS. The observation that many LOF variants are energetically destabilized and prone to protein misfolding motivates the idea of developing targeted therapies that assist in folding of destabilized KCNQ1 variants and/or rescue trafficking-defective variants to reach the plasma membrane. The development of such tailored therapies for varying classes of channel defects has become reality for the cystic fibrosis transmembrane regulator (CFTR) protein. Many patient-associated CFTR mutations are trafficking-defective, some of which have been shown to be rescuable by a pharmacological approach (78, 84). The structural models developed in this study provide a step toward realizing such tailored pharmacological approaches for KCNQ1.

### Conclusion

The structural models of the human KCNQ1 channel in closed and open states were developed by comparative modeling based on the high-grade template structure from *X. leavis* KCNQ1 and other experimental template structures that together provided nearly complete coverage of all channel regions (S0-S6). The KCNQ1 models are experimentally validated and offer improved completeness and accuracy over earlier KCNQ1 Rosetta models. MD simulations demonstrated the models’ capability to describe experimentally established channel properties such as state-dependent VSD gating charge interactions and pore conformations, PIP2 binding, and VSD-PD coupling interactions. Rosetta energy calculations on KCNQ1 models successfully predicted energetic destabilizations in folding-defective KCNQ1 LOF mutants and distinguished them from mutants with normal phenotype. The results in this work highlight the utility of the KCNQ1 structural models in addressing open questions about KCNQ1 structure-function mechanisms and assisting in the diagnosis of disease-causative KCNQ1 mutations towards the development of new therapeutic approaches.

## Materials and Methods

### Homology modeling of KCNQ1 RC state and AO state structures

Structural models of human KCNQ1 (helix S0-S6, residues 100 - 369) in the RC and AO state were generated by homology modeling using the protein structure prediction software Rosetta (version 3.8) (85). The *X. leavis* KCNQ1 cryo-EM structure (PDB 5VMS) (27) provided a template for the VSD in the AO model and a template for the pore domain in the RC model. Additional templates for the open pore conformation in AO state KCNQ1 were derived from the crystal structure of rat K_V_1.2-2.1 channel (PDB 2R9R) (31). Templates for the VSD in RC state KCNQ1 were obtained from related VSD structures: the resting VSD in *C. intestinalis* voltage-sensing phosphatase (Ci-VSP) (PDB 4G7Y) (28), VSD2 in *A. thaliana* two pore calcium channel protein 1 (TPC1) (PDB 5DQQ) (29), and the resting VSD conformation C3 in a model of the Shaker K channel (32). Hybridization and regularization of multiple templates was accomplished with the Rosetta comparative modeling (RosettaCM) protocol (86) guided by the RosettaMembrane energy function (87, 88). Missing loop regions and connecting residues between the VSD and pore domain were modeled *de novo* through fragment insertion. Template threading in RosettaCM was guided by multiple sequence alignments created with ClustalW (89) (see **Fig. S1** and **S2**). Fragment libraries were created with the Rosetta fragment picker (90) incorporating PSIPRED (91) secondary structure prediction. Transmembrane helix regions were predicted by OCTOPUS (92) and used to impose membrane-specific Rosetta energy terms on residues within the theoretical membrane bilayer. During homology modeling, C4 symmetry of the KCNQ1 tetramer was enforced by means of Rosetta symmetry definition files as described in DiMaio et al. (93). Amino acid side chain positions were optimized by a simulated annealing protocol, referred to as rotamer packing in Rosetta, and models were refined in internal and cartesian coordinate space by gradient-based minimization while applying a low harmonic restraint to the initial Cα-atom coordinates. Models were clustered based on Cα-atom RMSD, and the top-scoring models of the ten largest clusters as well as another 10-20 models with low Rosetta score and valid helix and loop conformations were selected as final ensemble. Model quality was additionally checked by MolProbity (34) analysis (see supporting **Tab. S1**), and the model with the lowest Molprobity score was considered the final model.

### MD simulations of KCNQ1 RC and AO models

MD simulations of KCNQ1 RC and AO models were performed in explicit phospholipid membranes at 310 K using Amber16 (41) and the Lipid17 force field *(Gould, I.R., Skjevik A.A., Dickson, C.J., Madej, B.D., Walker, R.C., “Lipid17: A Comprehensive AMBER Force Field for the Simulation of Zwitterionic and Anionic Lipids”, 2018, manuscript in preparation)*. Representative starting conformations for MD were selected from the ensemble of KCNQ1 homology models and by hierarchical full-linkage clustering.

Models were aligned to the membrane normal using the PPM webserver (94) and embedded into bilayers of POPC (palmitoyloleoyl-phosphatidylcholine) and PIP2 (phosphatidyl-4,5-bisphosphate) (~280 lipids per bilayer) using the membrane builder tool of the CHARMM-GUI website (95). A TIP3P water layer with 25 Å thickness containing 150 mM of KCl was added on either side of the membrane. In addition, four K^+^ ions were placed in the channel selectivity filter at positions inferred from the X-ray structure coordinates of PDB 2R9R. Bilayers contained 10 mol% of PIP2 in the inner leaflet which comprised equal numbers of C4-PO4- and C5-PO4-mono-protonated PIP2 molecules with stearoyl and arachidonoyl conjugations at the sn-1 and sn-2 position. The geometry of PIP2 was optimized with Gaussian 09 (Gaussian, Inc., Wallingford CT) on the B3LYP/6-31G* level of theory, and assignment of Amber atom types and calculation of RESP charges was done with Antechamber (96). Bond and angle parameters of the protonated C4-PO4 or C5-PO4 group in PIP2 were adjusted to values previously reported for phosphorylated amino acids (97) to avoid simulation instabilities. SHAKE (98) bond length constraints were applied to all bonds involving hydrogen. Nonbonded interactions were evaluated with a 10 Å cutoff, and electrostatic interactions were calculated by the particle-mesh Ewald method (99).

Each MD system was first minimized for 15,000 steps using steepest descent followed by 15,000 steps of conjugate gradient minimization. With protein and ions restrained to their initial coordinates, the lipid and water were heated to 50 K over 1000 steps with a step size of 1 fs in the NVT ensemble using Langevin dynamics with a rapid collision frequency of 10,000 ps^-1^. The system was then heated to 100 K over 50,000 steps with a collision frequency of 1000 ps^-1^ and finally to 310 K over 200,000 steps and a collision frequency of 100 ps^-1^. After changing to the NPT ensemble, restraints on ions were gradually removed over 500 ps and the system was equilibrated for another 5 ns at 310 K with weak positional restraints (with a force constant of 1 kcal mol^-1^ Å^-2^) applied to protein Cα atoms. The protein restraints were then gradually removed over 10 ns, and production MD was conducted for 400 ns using a step size of 2 fs, constant pressure periodic boundary conditions, anisotropic pressure scaling and Langevin dynamics. Four independent simulations were carried out for the RC as well as AO channel model yielding 2 x 1.6 μs of MD data.

### Analysis of KCNQ1 MD simulations

Analysis of MD trajectories with CPPTRAJ (version 18.0) (100) included calculation of Cα-atom root-mean-square deviations (Cα-RMSD) and root-mean-square fluctuations (Cα-RMSF), enumeration of protein-protein and protein-lipid hydrogen bonds and KCNQ1 residue contacts as well as principal component analysis. Residue contact numbers were calculated by counting the number of heteroatoms within a 4 Å distance of any heteroatom in a given KCNQ1 residue averaged over the last 300 ns of production MD as well as over all four RC and AO state MD simulations, respectively. For calculating intra-subunit contact numbers, only residues within the same chain were used whereas inter-subunit contact numbers were calculated considering only residues located on two different, directly adjacent chains. Intra- and inter-subunit contacts numbers were averaged over all four chains (i.e. A – D) or pairs of adjacent chains (i.e. A+B, B+C, C+D, A+D) in the KCNQ1 tetramer, respectively. Pairs of diagonally opposite chains (i.e. A+C and B+D) were not included in the analysis because they make no contacts between the VSD and PD. Furthermore, only long-range contacts (|i-j| > 5 residues) were used in the analysis of intra-subunit interactions.

Measurement of the channel pore radius was carried out with the HOLE program (101) and using snapshots of KCNQ1 taken at 1 ns intervals during the last 300 ns of MD.

### Stability calculations of KCNQ1 VSD variants

Free energy changes in 50 functionally characterized KCNQ1 variants (15) with mutations in the VSD were modeled using the Rosetta Flex ddG protocol (60) and the RosettaMembrane all-atom energy function (88). In short, the Flex ddG protocol models mutation-induced conformational and energetic changes by a series of “backrub” moves (15,000 steps in this study) of the protein backbone together with side-chain repacking within an 8 Å shell around the mutation site, followed by global minimization of all protein backbone and side-chain torsion angles. Minimization is performed with harmonic Cα-atom pair distance restraints to avoid any large structural deviations from the input model. A mutation was modeled in each of the four channel subunits simultaneously by enforcing C4 symmetry by means of Rosetta symmetry definition files. 50 independent trajectories were carried out for each mutant model and the WT model, and the Rosetta energy change (ΔΔG) was calculated as average score difference between the three top-scoring mutant and WT models. Stability calculations were repeated on ten different input models of the RC as well as AO state taken from the selected ensemble of KCNQ1 homology models to calculate the average ΔΔG value (± S.E.M.) for each KCNQ1 variant.

## Acknowledgements

This work was supported by NIH grants R01 HL122010 and R01 GM080403. GK was supported by fellowships from the German Research Foundation (KU 3510/1-1) and the American Heart Association (18POST34080422). This work was conducted using the resources of the Advanced Computing Center for Research and Education (ACCRE) at Vanderbilt University.

## Author contributions

JM, CRS, ALG, CGV and GK conceived this work and directed the approaches used. Model building and validation of the RC model was carried out by AMD with final model refinement done by GK. Model building and validation of the AO model was conducted by GK. MD simulations of KCNQ1 models were performed and analyzed by EFM under the supervision of GK. Rosetta ΔΔG calculations of KCNQ1 models were conducted and analyzed by KRB with guidance from GK. Application of the Rosetta ΔΔG method to membrane proteins was developed by HW, AMD and GK. ALG and CGV assisted with the interpretation of the KCNQ1 mutant channel phenotypes. GK wrote the paper with inputs from all authors. The final manuscript has been approved by all authors.

## Supporting Information Legends

**S1 Appendix**. Supporting tables and figures

**S1 PDB**. Coordinates of the Rosetta KCNQ1 AO model in PDB format

**S2 PDB**. Coordinates of the Rosetta KCNQ1 RC model in PDB format

## References

1. Abbott GW. Biology of the KCNQ1 Potassium Channel. New Journal of Science. 2014;2014:26.

2. Bezanilla F, Stefani E. Voltage-dependent gating of ionic channels. Annu Rev Biophys Biomol Struct. 1994;23:819–46.

3. Jensen MO, Jogini V, Borhani DW, Leffler AE, Dror RO, Shaw DE. Mechanism of voltage gating in potassium channels. Science. 2012;336(6078):229–33.

4. Bezanilla F. The voltage sensor in voltage-dependent ion channels. Physiol Rev. 2000;80(2):555–92.

5. Wu D, Delaloye K, Zaydman MA, Nekouzadeh A, Rudy Y, Cui J. State-dependent electrostatic interactions of S4 arginines with E1 in S2 during K_V_7.1 activation. J Gen Physiol. 2010;135(6):595–606.

6. Zaydman MA, Kasimova MA, McFarland K, Beller Z, Hou P, Kinser HE, et al. Domain-domain interactions determine the gating, permeation, pharmacology, and subunit modulation of the IKs ion channel. eLife. 2014;3:e03606–e.

7. Hou P, Eldstrom J, Shi J, Zhong L, McFarland K, Gao Y, et al. Inactivation of KCNQ1 potassium channels reveals dynamic coupling between voltage sensing and pore opening. Nat Commun. 2017;8(1):1730.

8. Barhanin J, Lesage F, Guillemare E, Fink M, Lazdunski M, Romey G. K(v)LQT1 and IsK (minK) proteins associate to form the I-Ks cardiac potassium current. Nature. 1996;384(6604):78–80.

9. Sanguinetti MC, Curran ME, Zou A, Shen J, Spector PS, Atkinson DL, et al. Coassembly of K(v)LQT1 and minK (IsK) proteins to form cardiac I-Ks potassium channel. Nature. 1996;384(6604):80–3.

10. Hedley PL, Jørgensen P, Schlamowitz S, Wangari R, Moolman-Smook J, Brink PA, et al. The genetic basis of long QT and short QT syndromes: a mutation update. Hum Mutat. 2009;30:1486–511.

11. Wu J, Ding WG, Horie M. Molecular pathogenesis of long QT syndrome type 1. J Arrhythm. 2016;32(5):381–8.

12. Eldstrom J, Wang Z, Werry D, Wong N, Fedida D. Microscopic mechanisms for long QT syndrome type 1 revealed by single-channel analysis of I(Ks) with S3 domain mutations in KCNQ1. Heart Rhythm. 2015;12:386–94.

13. Vanoye CG, Desai RR, Fabre KL, Gallagher SL, Potet F, DeKeyser JM, et al. High-Throughput Functional Evaluation of KCNQ1 Decrypts Variants of Unknown Significance. (2574-8300 (Electronic)).

14. Li B, Mendenhall JL, Kroncke BM, Taylor KC, Huang H, Smith DK, et al. Predicting the Functional Impact of KCNQ1 Variants of Unknown Significance. Circ Cardiovasc Genet. 2017;10(5).

15. Huang H, Kuenze G, Smith JA, Taylor KC, Duran AM, Hadziselimovic A, et al. Mechanisms of KCNQ1 channel dysfunction in long QT syndrome involving voltage sensor domain mutations. Sci Adv. 2018;4(3):eaar2631.

16. Smith JA, Vanoye CG, George AL, Meiler J, Sanders CR. Structural Models for the KCNQ1 Voltage-Gated Potassium Channel. Biochemistry. 2007;46(49):14141–52.

17. Kang C, Tian C, Sonnichsen FD, Smith JA, Meiler J, George AL, Jr., et al. Structure of KCNE1 and implications for how it modulates the KCNQ1 potassium channel. Biochemistry. 2008;47(31):7999–8006.

18. Chung DY, Chan PJ, Bankston JR, Yang L, Liu G, Marx SO, et al. Location of KCNE1 relative to KCNQ1 in the I(KS) potassium channel by disulfide cross-linking of substituted cysteines. Proc Natl Acad Sci U S A. 2009;106:743–8.

19. Chan PJ, Osteen JD, Xiong D, Bohnen MS, Doshi D, Sampson KJ, et al. Characterization of KCNQ1 atrial fibrillation mutations reveals distinct dependence on KCNE1. J Gen Physiol. 2012;139:135–44.

20. Nakajo K, Kubo Y. Steric hindrance between S4 and S5 of the KCNQ1/KCNE1 channel hampers pore opening. Nat Commun. 2014;5:4100-.

21. Li P, Liu H, Lai C, Sun P, Zeng W, Wu F, et al. Differential modulations of KCNQ1 by auxiliary proteins KCNE1 and KCNE2. Sci Rep. 2014;4:4973.

22. Kroncke BM, Van Horn WD, Smith J, Kang C, Welch RC, Song Y, et al. Structural basis for KCNE3 modulation of potassium recycling in epithelia. Sci Adv. 2016;2:e1501228–e.

23. Zaydman MA, Silva JR, Delaloye K, Li Y, Liang H, Larsson HP, et al. K_V_7.1 ion channels require a lipid to couple voltage sensing to pore opening. Proc Natl Acad Sci U S A. 2013;110:13180–5.

24. Eckey K, Wrobel E, Strutz-Seebohm N, Pott L, Schmitt N, Seebohm G. Novel K_V_7.1-phosphatidylinositol 4,5-bisphosphate interaction sites uncovered by charge neutralization scanning. J Biol Chem. 2014;289(33):22749–58.

25. Restier L, Cheng L, Sanguinetti MC. Mechanisms by which atrial fibrillation-associated mutations in the S1 domain of KCNQ1 slow deactivation of IKs channels. J Physiol. 2008;586:4179–91.

26. Labro AJ, Boulet IR, Choveau FS, Mayeur E, Bruyns T, Loussouarn G, et al. The S4-S5 linker of KCNQ1 channels forms a structural scaffold with the S6 segment controlling gate closure. J Biol Chem. 2011;286(1):717–25.

27. Sun J, MacKinnon R. Cryo-EM Structure of a KCNQ1/CaM Complex Reveals Insights into Congenital Long QT Syndrome. Cell. 2017;169:1042–50.e9.

28. Li Q, Wanderling S, Paduch M, Medovoy D, Singharoy A, McGreevy R, et al. Structural mechanism of voltage-dependent gating in an isolated voltage-sensing domain. Nat Struct Mol Biol. 2014;21:244–52.

29. Kintzer AF, Stroud RM. Structure, inhibition and regulation of two-pore channel TPC1 from Arabidopsis thaliana. Nature. 2016;531:258–62.

30. Yarov-Yarovoy V, Baker D, Catterall WA. Voltage sensor conformations in the open and closed states in ROSETTA structural models of K(+) channels. Proc Natl Acad Sci U S A. 2006;103(19):7292–7.

31. Long SB, Tao X, Campbell EB, MacKinnon R. Atomic structure of a voltage-dependent K+ channel in a lipid membrane-like environment. Nature. 2007;450(7168):376–82.

32. Henrion U, Renhorn J, Borjesson SI, Nelson EM, Schwaiger CS, Bjelkmar P, et al. Tracking a complete voltage-sensor cycle with metal-ion bridges. Proc Natl Acad Sci U S A. 2012;109(22):8552–7.

33. Laskowski RA, MacArthur MW, Moss DS, Thornton JM. PROCHECK: a program to check the stereochemical quality of protein structures. Journal of Applied Crystallography. 1993;26(2):283–91.

34. Davis IW, Leaver-Fay A, Chen VB, Block JN, Kapral GJ, Wang X, et al. MolProbity: all-atom contacts and structure validation for proteins and nucleic acids. Nucleic Acids Res. 2007;35(Web Server issue):W375–83.

35. Seebohm G, Strutz-Seebohm N, Ureche ON, Baltaev R, Lampert A, Kornichuk G, et al. Differential roles of S6 domain hinges in the gating of KCNQ potassium channels. Biophys J. 2006;90(6):2235–44.

36. Glauner KS, Mannuzzu LM, Gandhi CS, Isacoff EY. Spectroscopic mapping of voltage sensor movement in the Shaker potassium channel. Nature. 1999;402(6763):813–7.

37. Panaghie G, Abbott GW. The role of S4 charges in voltage-dependent and voltage-independent KCNQ1 potassium channel complexes. J Gen Physiol. 2007;129(2):121–33.

38. Xu Y, Wang Y, Meng X-Y, Zhang M, Jiang M, Cui M, et al. Building KCNQ1/KCNE1 channel models and probing their interactions by molecular-dynamics simulations. Biophys J. 2013;105:2461–73.

39. Yang T, Smith JA, Leake BF, Sanders CR, Meiler J, Roden DM. An allosteric mechanism for drug block of the human cardiac potassium channel KCNQ1. Mol Pharmacol. 2013;83(2):481–9.

40. Chen L, Zhang Q, Qiu Y, Li Z, Chen Z, Jiang H, et al. Migration of PIP2 lipids on voltage-gated potassium channel surface influences channel deactivation. Sci Rep. 2015;5:15079.

41. Case DA, Betz RM, Cerutti DS, Cheatham lii TE, Darden TA, Duke RE, et al. AMBER 2016. San Francisco: University of California; 2016.

42. Suh BC, Hille B. Regulation of ion channels by phosphatidylinositol 4,5-bisphosphate. Curr Opin Neurobiol. 2005;15(3):370–8.

43. Dickson CJ, Madej BD, Skjevik AA, Betz RM, Teigen K, Gould IR, et al. Lipid14: The Amber Lipid Force Field. J Chem Theory Comput. 2014;10(2):865–79.

44. Kasimova MA, Zaydman MA, Cui J, Tarek M. PIP2-dependent coupling is prominent in K_V_7.1 due to weakened interactions between S4-S5 and S6. Sci Rep. 2015;5:7474-.

45. Itoh H, Shimizu W, Hayashi K, Yamagata K, Sakaguchi T, Ohno S, et al. Long QT syndrome with compound mutations is associated with a more severe phenotype: a Japanese multicenter study. Heart Rhythm. 2010;7(10):1411–8.

46. Wisten A, Bostrom IM, Morner S, Stattin EL. Mutation analysis of cases of sudden unexplained death, 15 years after death: prompt genetic evaluation after resuscitation can save future lives. Resuscitation. 2012;83(10):1229–34.

47. Wang Q, Curran ME, Splawski I, Burn TC, Millholland JM, VanRaay TJ, et al. Positional cloning of a novel potassium channel gene: K_V_LQT1 mutations cause cardiac arrhythmias. Nat Genet. 1996;12:17–23.

48. Kapplinger JD, Tester DJ, Salisbury BA, Carr JL, Harris-Kerr C, Pollevick GD, et al. Spectrum and prevalence of mutations from the first 2,500 consecutive unrelated patients referred for the FAMILION long QT syndrome genetic test. Heart Rhythm. 2009;6(9):1297–303.

49. Napolitano C, Priori SG, Schwartz PJ, Bloise R, Ronchetti E, Nastoli J, et al. Genetic testing in the long QT syndrome: development and validation of an efficient approach to genotyping in clinical practice. JAMA. 2005;294(23):2975–80.

50. Millat G, Chevalier P, Restier-Miron L, Da Costa A, Bouvagnet P, Kugener B, et al. Spectrum of pathogenic mutations and associated polymorphisms in a cohort of 44 unrelated patients with long QT syndrome. Clin Genet. 2006;70(3):214–27.

51. Kubota T, Shimizu W, Kamakura S, Horie M. Hypokalemia-induced long QT syndrome with an underlying novel missense mutation in S4-S5 linker of KCNQ1. J Cardiovasc Electrophysiol. 2000; 11(9):1048–54.

52. Choi G, Kopplin LJ, Tester DJ, Will ML, Haglund CM, Ackerman MJ. Spectrum and frequency of cardiac channel defects in swimming-triggered arrhythmia syndromes. Circulation. 2004;110(15):2119–24.

53. Boulet IR, Labro AJ, Raes AL, Snyders DJ. Role of the S6 C-terminus in KCNQ1 channel gating. J Physiol. 2007;585(Pt 2):325–37.

54. Choveau FS, Rodriguez N, Abderemane Ali F, Labro AJ, Rose T, Dahimene S, et al. KCNQ1 channels voltage dependence through a voltage-dependent binding of the S4-S5 linker to the pore domain. J Biol Chem. 2011;286(1):707–16.

55. Ma LJ, Ohmert I, Vardanyan V. Allosteric features of KCNQ1 gating revealed by alanine scanning mutagenesis. Biophys J. 2011;100(4):885–94.

56. Lu Z, Klem AM, Ramu Y. Ion conduction pore is conserved among potassium channels. Nature. 2001;413(6858):809–13.

57. Long SB, Campbell EB, Mackinnon R. Voltage sensor of K_V_1.2: structural basis of electromechanical coupling. Science. 2005;309(5736):903–8.

58. Lu Z, Klem AM, Ramu Y. Coupling between voltage sensors and activation gate in voltage-gated K+ channels. J Gen Physiol. 2002;120(5):663–76.

59. Silva JR, Pan H, Wu D, Nekouzadeh A, Decker KF, Cui J, et al. A multiscale model linking ion-channel molecular dynamics and electrostatics to the cardiac action potential. Proc Natl Acad Sci U S A. 2009; 106(27):11102–6.

60. Barlow KA, S Oc, Thompson S, Suresh P, Lucas JE, Heinonen M, et al. Flex ddG: Rosetta Ensemble-Based Estimation of Changes in Protein-Protein Binding Affinity upon Mutation. J Phys Chem B. 2018;122(21):5389–99.

61. Davis IW, Arendall WB, 3rd, Richardson DC, Richardson JS. The backrub motion: how protein backbone shrugs when a sidechain dances. Structure. 2006;14(2):265–74.

62. Alford RF, Koehler Leman J, Weitzner BD, Duran AM, Tilley DC, Elazar A, et al. An Integrated Framework Advancing Membrane Protein Modeling and Design. PLoS Comput Biol. 2015;11(9):e1004398.

63. Tinel N, Diochot S, Borsotto M, Lazdunski M, Barhanin J. KCNE2 confers background current characteristics to the cardiac KCNQ1 potassium channel. EMBO J. 2000;19(23):6326–30.

64. Schroeder BC, Waldegger S, Fehr S, Bleich M, Warth R, Greger R, et al. A constitutively open potassium channel formed by KCNQ1 and KCNE3. Nature. 2000;403(6766):196–9.

65. Nakajo K, Kubo Y. KCNQ1 channel modulation by KCNE proteins via the voltage-sensing domain. J Physiol. 2015;593:2617–25.

66. Tapper AR, George AL. Location and orientation of minK within the I-Ks potassium channel complex. Journal of Biological Chemistry. 2001;276(41):38249–54.

67. Xu X, Jiang M, Hsu K-L, Zhang M, Tseng G-N. KCNQ1 and KCNE1 in the IKs channel complex make state-dependent contacts in their extracellular domains. J Gen Physiol. 2008;131:589–603.

68. Wang YH, Jiang M, Xu XL, Hsu K-L, Zhang M, Tseng G-N. Gating-related molecular motions in the extracellular domain of the IKs channel: implications for IKs channelopathy. J Membr Biol. 2011;239:137–56.

69. Strutz-Seebohm N, Pusch M, Wolf S, Stoll R, Tapken D, Gerwert K, et al. Structural basis of slow activation gating in the cardiac I Ks channel complex. Cell Physiol Biochem. 2011;27:443–52.

70. Wang Y, Zhang M, Xu Y, Jiang M, Zankov DP, Cui M, et al. Probing the structural basis for differential KCNQ1 modulation by KCNE1 and KCNE2. J Gen Physiol. 2012;140(6):653–69.

71. Barro-Soria R, Perez ME, Larsson HP. KCNE3 acts by promoting voltage sensor activation in KCNQ1. Proc Natl Acad Sci U S A. 2015;112(52):E7286–92.

72. Nakajo K, Kubo Y. KCNE1 and KCNE3 stabilize and/or slow voltage sensing S4 segment of KCNQ1 channel. J Gen Physiol. 2007;130(3):269–81.

73. Barro-Soria R, Ramentol R, Liin SI, Perez ME, Kass RS, Larsson HP. KCNE1 and KCNE3 modulate KCNQ1 channels by affecting different gating transitions. Proc Natl Acad Sci U S A. 2017;114(35):E7367–E76.

74. Stefl S, Nishi H, Petukh M, Panchenko AR, Alexov E. Molecular mechanisms of disease-causing missense mutations. J Mol Biol. 2013;425(21):3919–36.

75. Kroncke BM, Vanoye CG, Meiler J, George AL, Jr., Sanders CR. Personalized biochemistry and biophysics. Biochemistry. 2015;54(16):2551–9.

76. Berliner N, Teyra J, Colak R, Garcia Lopez S, Kim PM. Combining structural modeling with ensemble machine learning to accurately predict protein fold stability and binding affinity effects upon mutation. PLoS One. 2014;9(9):e107353.

77. Kroncke BM, Duran AM, Mendenhall JL, Meiler J, Blume JD, Sanders CR. Documentation of an Imperative To Improve Methods for Predicting Membrane Protein Stability. Biochemistry. 2016;55(36):5002–9.

78. Marinko JT, Huang H, Penn WD, Capra JA, Schlebach JP, Sanders CR. Folding and Misfolding of Human Membrane Proteins in Health and Disease: From Single Molecules to Cellular Proteostasis. Chem Rev. 2019.

79. Smith JL, Anderson CL, Burgess DE, Elayi CS, January CT, Delisle BP. Molecular pathogenesis of long QT syndrome type 2. J Arrhythm. 2016;32(5):373–80.

80. Schlebach JP, Narayan M, Alford C, Mittendorf KF, Carter BD, Li J, et al. Conformational Stability and Pathogenic Misfolding of the Integral Membrane Protein PMP22. J Am Chem Soc. 2015; 137(27):8758–68.

81. Farinha CM, Matos P, Amaral MD. Control of cystic fibrosis transmembrane conductance regulator membrane trafficking: not just from the endoplasmic reticulum to the Golgi. FEBS J. 2013;280(18):4396–406.

82. Swanton E, Holland A, High S, Woodman P. Disease-associated mutations cause premature oligomerization of myelin proteolipid protein in the endoplasmic reticulum. Proc Natl Acad Sci U S A. 2005; 102(12):4342–7.

83. Tester DJ, Will ML, Haglund CM, Ackerman MJ. Compendium of cardiac channel mutations in 541 consecutive unrelated patients referred for long QT syndrome genetic testing. Heart Rhythm. 2005;2(5):507–17.

84. Fajac I, De Boeck K. New horizons for cystic fibrosis treatment. Pharmacol Ther. 2017;170:205–11.

85. Bender BJ, Cisneros A, 3rd, Duran AM, Finn JA, Fu D, Lokits AD, et al. Protocols for Molecular Modeling with Rosetta3 and RosettaScripts. Biochemistry. 2016;55(34):4748–63.

86. Song Y, DiMaio F, Wang RY, Kim D, Miles C, Brunette T, et al. High-resolution comparative modeling with RosettaCM. Structure. 2013;21(10):1735–42.

87. Yarov-Yarovoy V, Schonbrun J, Baker D. Multipass membrane protein structure prediction using Rosetta. Proteins. 2006;62(4):1010–25.

88. Barth P, Schonbrun J, Baker D. Toward high-resolution prediction and design of transmembrane helical protein structures. Proc Natl Acad Sci U S A. 2007;104(40):15682–7.

89. Larkin MA, Blackshields G, Brown NP, Chenna R, McGettigan PA, McWilliam H, et al. Clustal W and Clustal X version 2.0. Bioinformatics. 2007;23(21):2947–8.

90. Gront D, Kulp DW, Vernon RM, Strauss CE, Baker D. Generalized fragment picking in Rosetta: design, protocols and applications. PLoS One. 2011;6(8):e23294.

91. Jones DT. Protein Secondary Structure Prediction Based on Position-specific Scoring Matrices. J Mol Biol. 1999;292(2):195–202.

92. Viklund H, Elofsson A. OCTOPUS: improving topology prediction by two-track ANN-based preference scores and an extended topological grammar. Bioinformatics. 2008;24(15):1662–8.

93. Dimaio F, Leaver-Fay A, Bradley P, Baker D, Andre I. Modeling symmetric macromolecular structures in rosetta3. PLoS One. 2011;6(6):e20450.

94. Lomize MA, Pogozheva ID, Joo H, Mosberg HI, Lomize AL. OPM database and PPM web server: resources for positioning of proteins in membranes. Nucleic Acids Res. 2012;40(Database issue):D370–6.

95. Wu EL, Cheng X, Jo S, Rui H, Song KC, Davila-Contreras EM, et al. CHARMM-GUI Membrane Builder toward realistic biological membrane simulations. J Comput Chem. 2014;35(27):1997–2004.

96. Wang J, Wang W, Kollman PA, Case DA. Automatic atom type and bond type perception in molecular mechanical calculations. J Mol Graph Model. 2006;25(2):247–60.

97. Homeyer N, Horn AH, Lanig H, Sticht H. AMBER force-field parameters for phosphorylated amino acids in different protonation states: phosphoserine, phosphothreonine, phosphotyrosine, and phosphohistidine. J Mol Model. 2006;12(3):281–9.

98. Ryckaert JP, Ciccotti G, Berendsen HJC. Numerical-Integration of Cartesian Equations of Motion of a System with Constraints - Molecular-Dynamics of N-Alkanes. J Comput Phys. 1977;23(3):327–41.

99. Darden T, York D, Pedersen L. Particle Mesh Ewald - an N. Log(N) Method for Ewald Sums in Large Systems. Journal of Chemical Physics. 1993;98(12):10089–92.

100. Roe DR, Cheatham TE, 3rd. PTRAJ and CPPTRAJ: Software for Processing and Analysis of Molecular Dynamics Trajectory Data. J Chem Theory Comput. 2013;9(7):3084–95.

101. Smart OS, Neduvelil JG, Wang X, Wallace BA, Sansom MS. HOLE: a program for the analysis of the pore dimensions of ion channel structural models. J Mol Graph. 1996;14(6):354–60, 76.

