## Supplemental Information for "Upgraded molecular models of the human KCNQ1 potassium channel"

### Supporting Tables

Table S1: Molprobity statistic of KCNQ1 AO and RC homology models

| | KCNQ1 AO state | KCNQ1 RC state |
|---|---|---|
| Molprobity score | 1.38 (97th percentile) | 1.71 (90th percentile) |
| Clash score | 3.09 (98th percentile) | 4.92 (94th percentile) |
| Ramachandran statistic | | |
| Favored regions (%) | 95.9 | 92.8 |
| Allowed regions (%) | 4.1 | 6.2 |
| Disallowed regions (%) | 0.0 | 1.0 |
| Rotamer statistic | | |
| Favored rotamers (%) | 100.0 | 99.5 |
| Poor rotamers (%) | 0.0 | 0.0 |
| Cβ deviations | 0 | 0 |
| Bad bonds (%) | 0.00 | 0.00 |
| Bad angles (%) | 0.03 | 0.03 |

Table S2: Comparison of Rosetta predicted stability changes (ΔΔG) with expression levels, trafficking efficiencies and peak current densities of KCNQ1 VSD mutants

| KCNQ1 variant | ClinVar Class | Functional Class* | Total expression (% WT)# | Cell surface expression (% WT) | Trafficking efficiency (% WT) | Peak current density (% WT) | ΔΔG RC model (REU) | ΔΔG AO model (REU) |
|---|---|---|---|---|---|---|---|---|
| V100I | neutral | VI | 101.1 ± 3.4 | 91.0 ± 4.4 | 112.4 ± 13.3 | 113.0 ± 14.0 | 0.7 ± 3.1 | 0.3 ± 2.2 |
| A102S | neutral | VI | 106.5 ± 4.9 | 95.8 ± 5.2 | 113.1 ± 16.8 | 282.0 ± 36.0 | -2.9 ± 1.7 | 0.2 ± 3.2 |
| T104I | VUS | IV | 89.0 ± 4.6 | 64.9 ± 5.1 | 90.7 ± 10.0 | 22.0 ± 4.0 | 2.2 ± 1.9 | 8.2 ± 2.7 |
| T104S | neutral | VI | 101.8 ± 5.0 | 93.2 ± 6.2 | 113.9 ± 11.8 | 82.0 ± 13.0 | 3.4 ± 1.3 | 5.5 ± 4.2 |
| H105L | LQTS | III | 97.6 ± 4.4 | 80.2 ± 7.5 | 101.6 ± 9.6 | 131.0 ± 17.0 | -5.5 ± 2.6 | 6.4 ± 2.1 |
| H105N | neutral | VI | 131.8 ± 6.2 | 105.6 ± 6.7 | 98.9 ± 4.9 | 72.0 ± 9.0 | -4.7 ± 1.8 | 0.0 ± 2.4 |
| H105Y | neutral | VI | 94.7 ± 2.6 | 78.4 ± 5.5 | 102.2 ± 7.5 | 151.0 ± 21.0 | 4.3 ± 4.3 | 6.8 ± 2.5 |
| V106I | neutral | VI | 121.4 ± 7.7 | 103.7 ± 4.2 | 106.7 ± 11.3 | 121.0 ± 18.0 | 1.2 ± 2.5 | 4.5 ± 1.4 |
| Q107H | VUS | II | 20.1 ± 2.5 | 5.6 ± 1.7 | 33.8 ± 8.5 | 36.0 ± 6.0 | 4.3 ± 3.7 | 12.6 ± 2.1 |
| R109L | VUS | III | 114.0 ± 2.8 | 91.9 ± 6.2 | 99.2 ± 4.5 | 223.0 ± 35.0 | 10.9 ± 2.6 | 10.3 ± 1.7 |
| V110I | LQTS | I | 114.1 ± 6.2 | 96.4 ± 5.5 | 104.8 ± 8.4 | 25.0 ± 5.0 | 3.2 ± 2.7 | 3.4 ± 2.2 |
| Y111C | LQTS | V | 17.6 ± 2.5 | 2.0 ± 1.1 | 13.5 ± 6.7 | -2.0 ± 3.0 | 19.5 ± 3.6 | 30.2 ± 2.3 |
| L114P | LQTS | V | 15.6 ± 2.1 | 2.0 ± 1.2 | 14.8 ± 7.7 | -1.0 ± 4.0 | 77.1 ± 4.0 | 70.5 ± 4.1 |
| E115G | LQTS | III | 18.5 ± 1.2 | 1.9 ± 1.0 | 12.6 ± 6.4 | -7.0 ± 3.0 | 27.6 ± 4.0 | 26.9 ± 4.6 |
| P117L | LQTS | V | 25.7 ± 2.5 | 5.5 ± 1.4 | 26.3 ± 5.9 | -5.0 ± 3.0 | 20.1 ± 5.9 | 24.2 ± 7.6 |
| T118S | neutral | III | 110.6 ± 1.5 | 100.7 ± 7.1 | 112.0 ± 4.7 | 76.0 ± 14.0 | 3.6 ± 1.9 | 0.5 ± 2.2 |
| C122Y | LQTS | I | 95.6 ±10.8 | 71.7 ± 4.0 | 66.8 ± 4.0 | -4.0 ± 3.0 | -3.5 ± 2.8 | -11.9 ± 2.9 |
| V124I | neutral | VI | 112.1 ±15.4 | 112.8 ± 3.5 | 79.8 ± 4.3 | 92.0 ± 12.0 | -1.5 ± 2.5 | -5.8 ± 1.8 |
| Y125D | VUS | V | 12.6 ± 1.4 | 2.1 ± 0.6 | 20.0 ± 5.4 | 1.0 ± 3.0 | 18.2 ± 1.7 | 27.6 ± 1.8 |

| | | | | | | | | |
|---|---|---|---|---|---|---|---|---|
| H126L | VUS | V | 32.4 ± 3.3 | 3.7 ± 0.9 | 4.2 ± 2.6 | 4.0 ± 3.0 | 4.1 ± 3.2 | 11.5 ± 6.0 |
| F127L | LQTS | VI | 107.6 ± 4.4 | 110.7 ± 4.1 | 85.1 ± 13.1 | 80.0 ± 13.0 | 0.3 ± 2.3 | -6.4 ± 2.2 |
| A128T | neutral | VI | 101.8 ± 8.5 | 112.8 ± 4.7 | 95.3 ± 19.3 | 74.0 ± 11.0 | -2.2 ± 2.1 | 0.7 ± 2.3 |
| V129I | VUS | VI | 120.7 ± 8.2 | 132.0 ± 6.7 | 91.1 ± 13.6 | 83.0 ± 16.0 | -4.4 ± 2.8 | 1.6 ± 2.3 |
| L131P | VUS | IV | 41.9 ± 4.8 | 45.4 ± 4.0 | 102.4 ± 20.9 | 6.0 ± 3.0 | 61.5 ± 2.5 | 76.0 ± 3.0 |
| I132L | LQTS | III | 109.9 ± 6.2 | 104.4 ± 6.5 | 90.7 ± 2.9 | 112.0 ± 23.0 | -4.9 ± 2.2 | -0.2 ± 1.7 |
| V133I | LQTS | VI | 110.9 ± 6.7 | 95.0 ± 3.9 | 85.0 ± 1.1 | 67.0 ± 16.0 | 5.3 ± 2.5 | 0.6 ± 2.8 |
| L134P | LQTS | I | 60.0 ± 3.2 | 99.6 ±11.4 | 177.7 ± 11.7 | 3.0 ± 2.0 | 63.7 ± 2.6 | 68.3 ± 4.1 |
| V135A | neutral | VI | 103.2 ± 4.7 | 92.3 ± 5.2 | 91.7 ± 1.3 | 78.0 ± 15.0 | 1.8 ± 3.0 | 5.4 ± 2.5 |
| V135I | neutral | VI | 99.1 ± 8.5 | 83.6 ± 7.6 | 92.7 ± 6.7 | 100.0 ± 11.0 | -7.0 ± 2.3 | -2.5 ± 2.3 |
| A149V | neutral | VI | 66.1 ± 2.7 | 73.4 ±10.2 | 127.2 ± 3.5 | 107.0 ± 34.0 | 16.1 ± 3.7 | 26.5 ± 2.3 |
| A150T | VUS | I | 117.4 ± 7.5 | 156.1 ±10.7 | 139.3 ± 1.1 | 58.0 ± 9.0 | 1.4 ± 2.1 | -4.4 ± 2.3 |
| A150V | neutral | I | 90.6 ± 5.5 | 96.8 ± 4.7 | 113.8 ± 9.7 | 52.0 ± 17.0 | 5.9 ± 1.6 | 3.4 ± 2.4 |
| E160K | LQTS | IV | 35.5 ± 3.0 | 15.8 ± 1.6 | 55.0 ± 3.5 | -4.0 ± 3.0 | 19.5 ± 4.2 | 16.3 ± 5.2 |
| T169M | VUS | I | 87.8 ± 4.5 | 86.5 ± 4.7 | 121.6 ± 5.2 | 65.0 ± 10.0 | -7.6 ± 1.9 | -4.4 ± 2.9 |
| R174C | LQTS | V | 14.0 ± 4.0 | 2.2 ± 1.2 | 17.2 ± 6.8 | 0.0 ± 2.0 | 40.0 ± 3.0 | 36.3 ± 3.5 |
| R174H | LQTS | V | 20.0 ± 2.7 | 4.2 ± 1.3 | 25.6 ± 6.2 | -1.0 ± 3.0 | 37.7 ± 2.0 | 37.1 ± 4.2 |
| R174L | LQTS | V | 17.5 ± 2.4 | 9.7 ± 1.8 | 68.0 ± 7.5 | 3.0 ± 3.0 | 27.0 ± 2.0 | 31.4 ± 4.1 |
| W176R | VUS | V | 10.8 ± 1.9 | 2.5 ± 1.7 | 24.1 ± 14.4 | 1.0 ± 2.0 | 14.7 ± 2.8 | 9.3 ± 3.6 |
| G179S | LQTS | V | 17.7 ± 3.2 | 4.1 ± 2.2 | 25.3 ± 12.8 | 10.0 ± 3.0 | -5.6 ± 1.9 | 2.6 ± 2.5 |
| G189A | VUS | V | 14.1 ± 0.8 | 2.4 ± 1.7 | 19.6 ± 13.3 | -5.0 ± 2.0 | 13.2 ± 1.6 | 21.6 ± 3.8 |
| R195P | VUS | V | 16.8 ± 2.8 | 2.0 ± 1.6 | 11.6 ± 8.6 | -1.0 ± 3.0 | 52.7 ± 1.1 | 56.8 ± 2.0 |
| K196T | VUS | IV | 61.6 ± 3.9 | 58.3 ± 2.8 | 116.9 ± 2.4 | 20.0 ± 5.0 | 1.0 ± 3.1 | 5.2 ± 3.4 |
| P197L | VUS | II | 68.1 ± 3.0 | 59.1 ± 3.7 | 107.3 ± 7.0 | 147.0 ± 23.0 | -11.6 ± 2.8 | -10.9 ± 2.6 |
| P197S | VUS | IV | 40.2 ± 4.4 | 21.2 ± 2.3 | 65.2 ± 0.3 | 17.0 ± 4.0 | 6.9 ± 1.4 | 10.7 ± 4.0 |
| V207M | LQTS | VI | 85.6 ± 4.4 | 70.8 ± 4.1 | 102.0 ± 2.7 | 88.0 ± 15.0 | 4.2 ± 4.3 | 4.9 ± 2.4 |
| K218E | VUS | I | 93.1 ± 5.3 | 101.8 ± 3.5 | 135.5 ± 7.8 | 46.0 ± 9.0 | 12.4 ± 1.7 | 8.4 ± 2.4 |
| I227L | VUS | I | 116.0 ± 7.5 | 142.9 ± 9.8 | 152.0 ± 6.1 | 22.0 ± 5.0 | -7.5 ± 2.2 | 4.4 ± 2.0 |
| Q234P | VUS | I | 116.4 ± 6.8 | 131.8 ± 1.5 | 140.1 ± 3.7 | 3.0 ± 4.0 | 82.1 ± 3.8 | 71.0 ± 5.0 |
| L236P | VUS | IV | 78.1 ± 4.7 | 47.0 ± 8.3 | 72.0 ± 7.3 | 3.0 ± 4.0 | 60.3 ± 4.8 | 65.8 ± 6.0 |
| L236R | VUS | IV | 56.1 ± 2.7 | 15.6 ± 3.5 | 33.5 ± 5.8 | 1.0 ± 4.0 | 13.9 ± 2.6 | 19.0 ± 3.2 |

*Functional classification of KCNQ1 variants according to reference (1):

**Class I:** Normal or higher expression and trafficking levels but low peak current density. Dysfunctional channel.

**Class II:** Lower expression levels but normal channel function for channel variants that traffic to the membrane.

**Class III:** Normal or higher expression and trafficking levels and peak current density but altered channel $V_{1/2}$ and/or deactivation rate.

**Class IV:** Defective in surface expression levels and electrophysiological channel properties.

**Class V:** Severe expression and/or trafficking defects. Current is so low that channel properties cannot be assessed.

**Class VI:** Have wildtype-like behavior. Normal or higher surface expression levels and channel properties.

[#] Experimental data are from reference (1). Values are reported as mean ± S.E.M.

### Supporting Figures

#### Figure S1: Multiple sequence alignment of human KCNQ1 for RC homology modeling

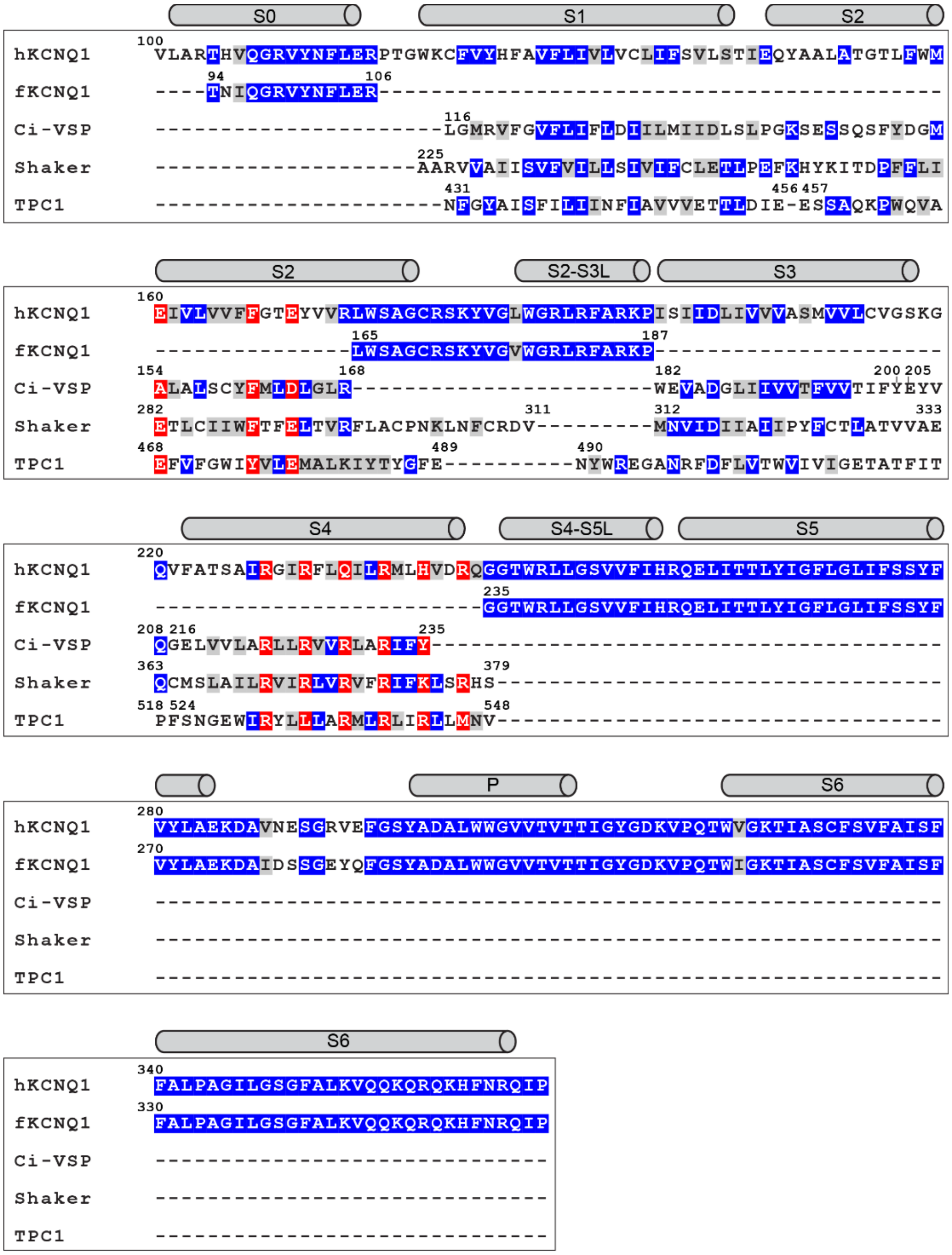

**Figure S1: Multiple sequence alignment of human KCNQ1 for RC homology modeling.** *C. intestinalis* voltage-sensing phosphatase (Ci-VSP) (PDB 4G7Y), VSD2 in *A. thaliana* two pore calcium channel protein 1 (TPC1) (PDB 5DQQ), and the resting VSD conformation C3 in a model of the Shaker K channel were used as structural templates for helix S1, S2, S3 and S4 of the VSD. The cryo-EM structure of *X. leavis* KCNQ1 (fKCNQ1) (PDB 5VMS) was used as template for helix S0, the S2-S3 linker and the pore domain. The alignment was created with MAMMOTH (2) and ClustalW (3) and manually adjusted to ensure that functionally conserved residues in S2, S3 and S4 are correctly aligned. Identical and similar residues are colored blue and gray, respectively. Residues at structurally conserved positions are highlighted in red. Predicted secondary structure regions are indicated above the sequence alignment as gray cylinders.

Figure S2: Multiple sequence alignment of human KCNQ1 for AO homology modeling

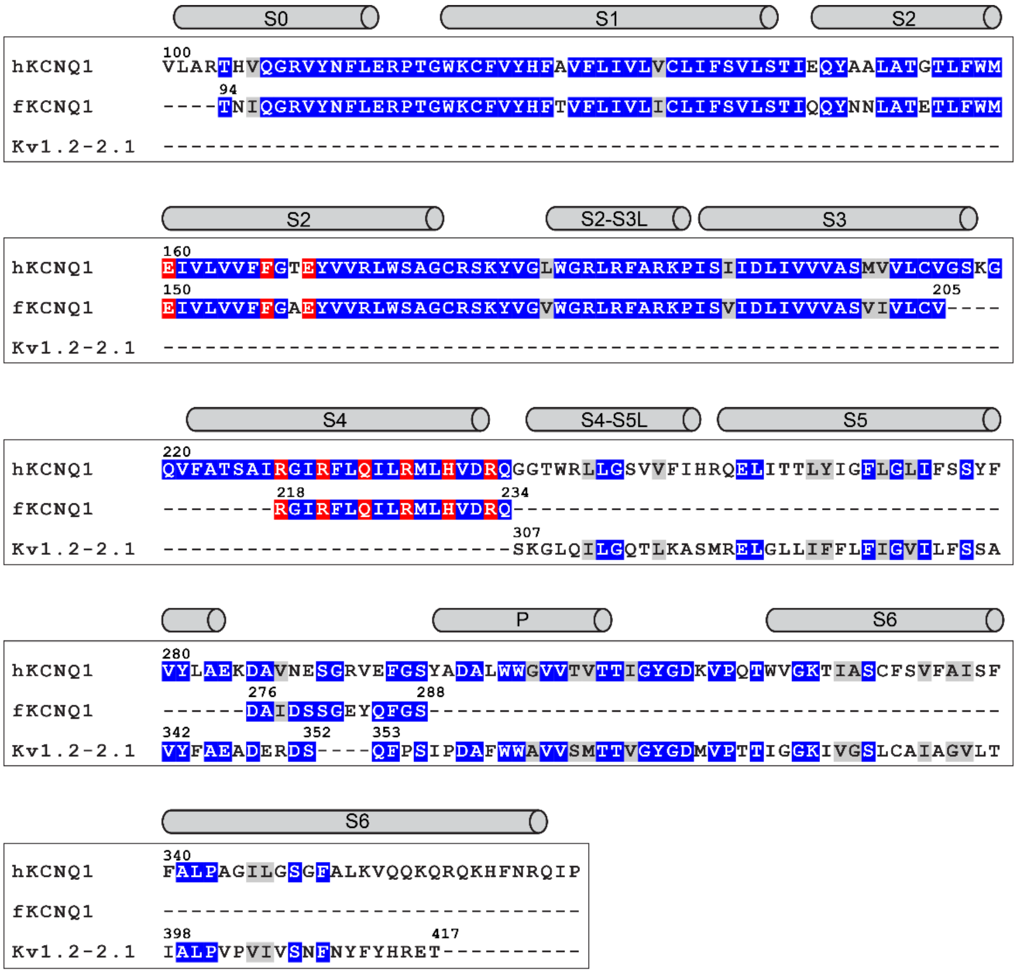

**Figure S2: Multiple sequence alignment of human KCNQ1 for AO homology modeling.** The cryo-EM structure of *X. leavis* KCNQ1 (fKCNQ1) (PDB 5VMS) and the X-ray structure of the rat chimeric $K_V$1.2-2.1 channel (PDB 2R9R) were used as structural templates. The alignment was created with MAMMOTH (2) and ClustalW (3) and manually adjusted to ensure functionally conserved residues in S2, S3 and S4 are correctly aligned. Identical and similar residues are colored blue and gray, respectively. Residues at structurally conserved positions are highlighted in red. Predicted secondary structure regions are indicated above the sequence alignment as gray cylinders.

Figure S3: Pore radius of KCNQ1 in MD simulations of RC and AO models

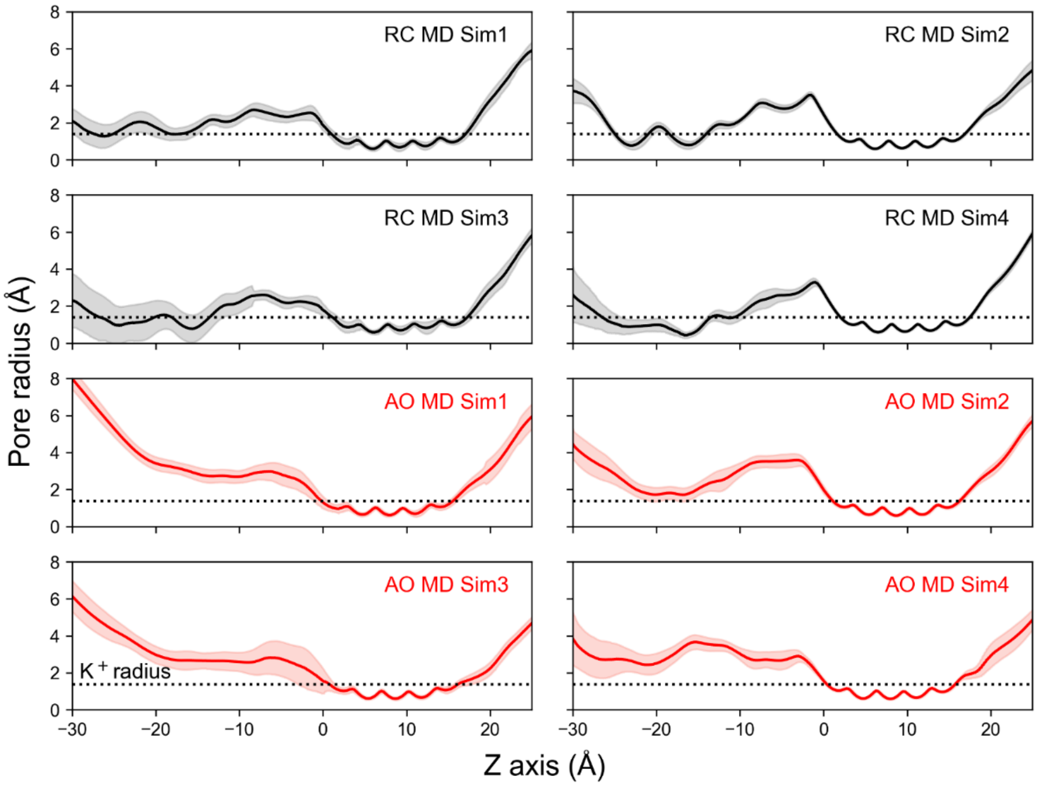

**Figure S3: Pore radius of KCNQ1 in MD simulations of RC and AO models.** Four MD simulations of the RC and AO state were conducted in this study. The average pore radius was calculated over the last 300 ns of production MD and is shown as solid line. Shaded areas correspond to one standard deviation. The approximate radius of a $K^+$ ion is indicated as dashed line. The region between 3Å – 14Å corresponds to the channel selectivity filter.

Figure S4: Backbone Cα-RMSD of KCNQ1 RC and AO state MD trajectories

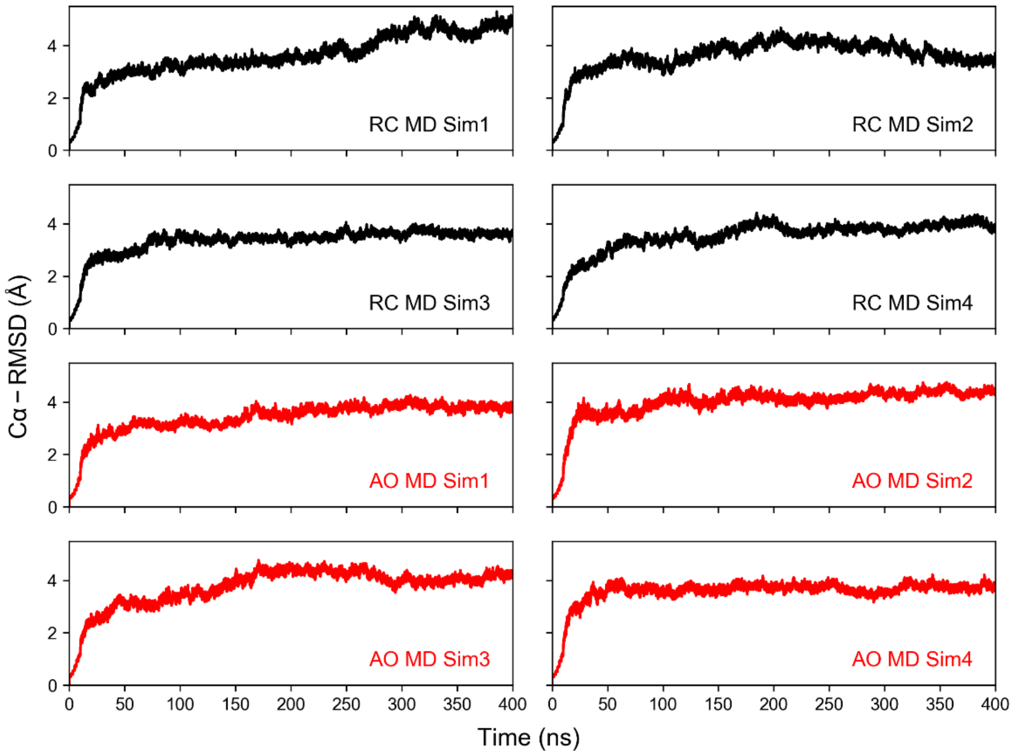

**Figure S4: Backbone Cα-RMSD of KCNQ1 RC and AO state MD trajectories.** Four MD simulations of the RC and AO state were conducted in this study. Each simulation started from a different model from the final ensemble of 20-30 Rosetta homology models and was conducted for 400 ns as described in Methods. The RMSD is displayed for the production period of MD and was calculated relative to the conformation after minimization and system heating.

Figure S5: Global domain motions in KCNQ1 channel models revealed by residue cross-correlation and principal component analysis

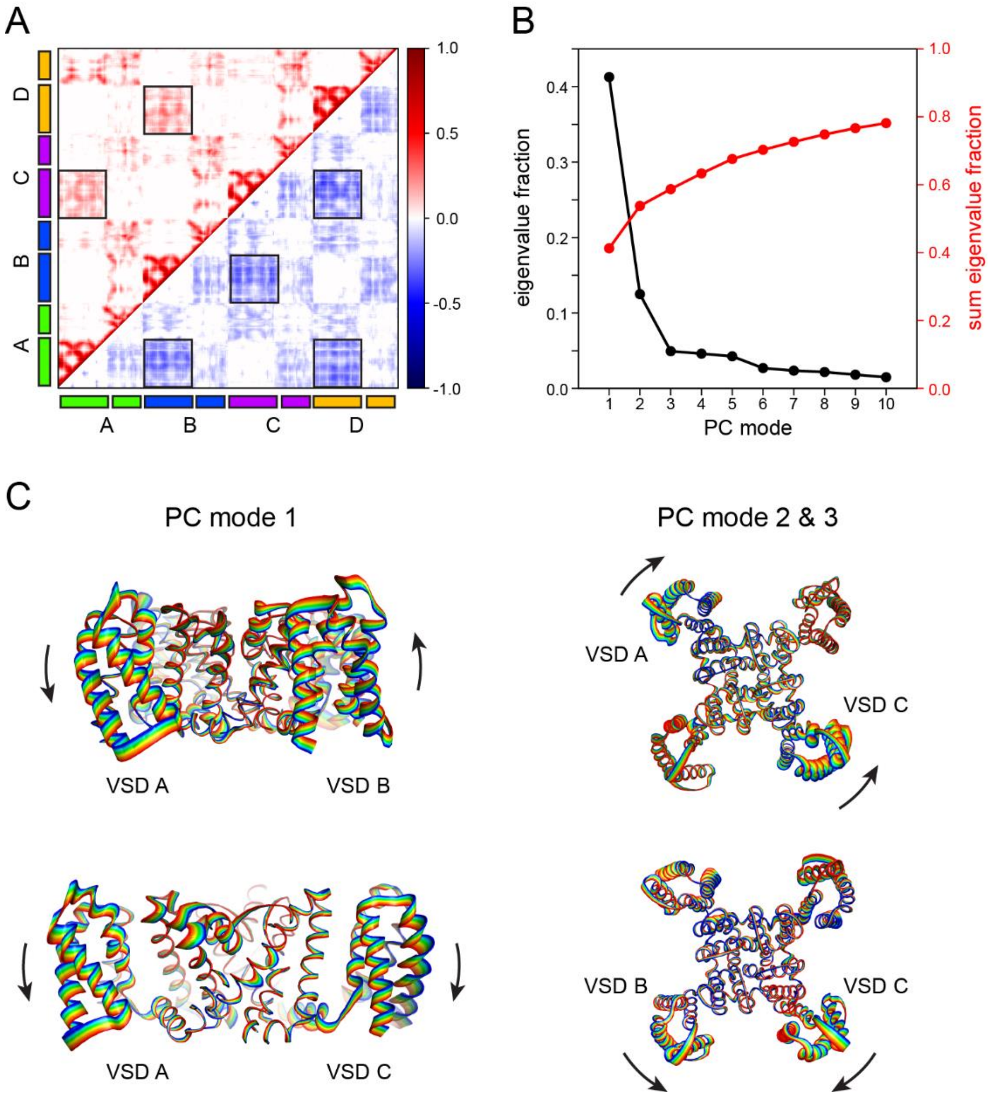

**Figure S5: Global domain motions in KCNQ1 channel models revealed by residue cross-correlation and principal component analysis.** (**A**) Average dynamic cross-correlation matrix of the KCNQ1 tetramer calculated from MD trajectories of the AO models. Positive residue correlations are plotted in the upper triangular matrix whereas negative correlations are mapped on the lower triangle. Regions in the cross-correlation matrix corresponding to correlations between VSDs are framed by black boxes. The approximate regions of the four channel subunits (labeled A – D) are indicated on the x- and y-axis. (**B**) Scree plot of the first ten principal components obtained by PCA of the KCNQ1 MD simulations. (**C**) Pseudo-trajectories along the first three PC modes. PC mode 1 corresponds to a VSD movement along the membrane normal with two VSDs on the same side of the channel tetramer moving anti-parallel and VSDs on opposite sides moving in a parallel fashion. PC modes 2 and 3 represent a swing movement of the VSDs within the membrane plane.

Figure S6: Location of mutation sites in the KCNQ1 VSD by variant class.

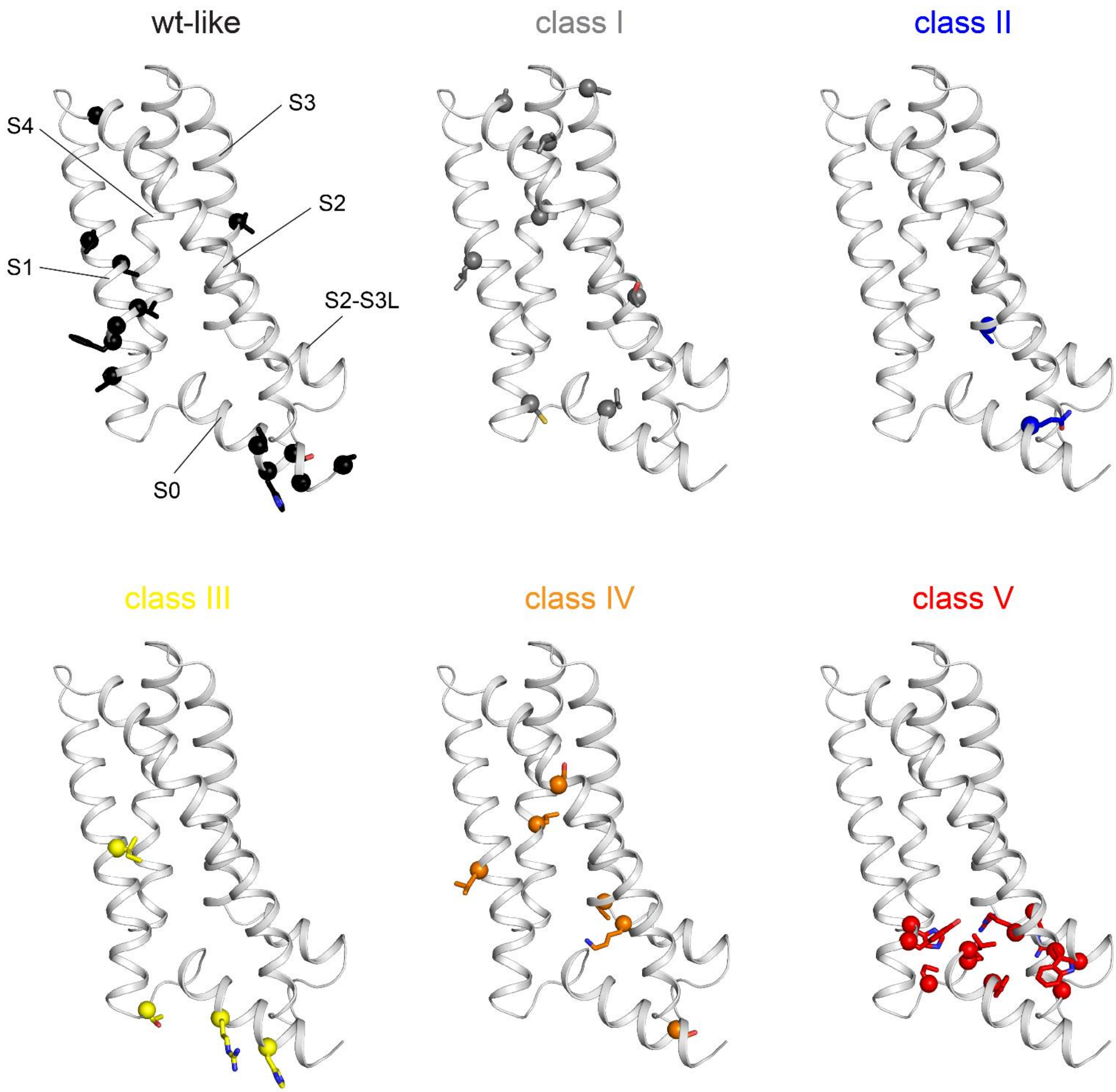

**Figure S6: Location of mutation sites in the KCNQ1 VSD by variant class.** The VSD is represented as ribbon with helical segments labeled. The backbone position of mutation sites is indicated by a sphere and the native amino acid residue is shown in sticks. The assignment of variants to these six classes can be found in **Tab. S2** and reference (1) and the class definition is given in the footnote of **Tab. S2**.

Figure S7: Predicted stability changes of KCNQ1 VSD variants calculated with the RC model

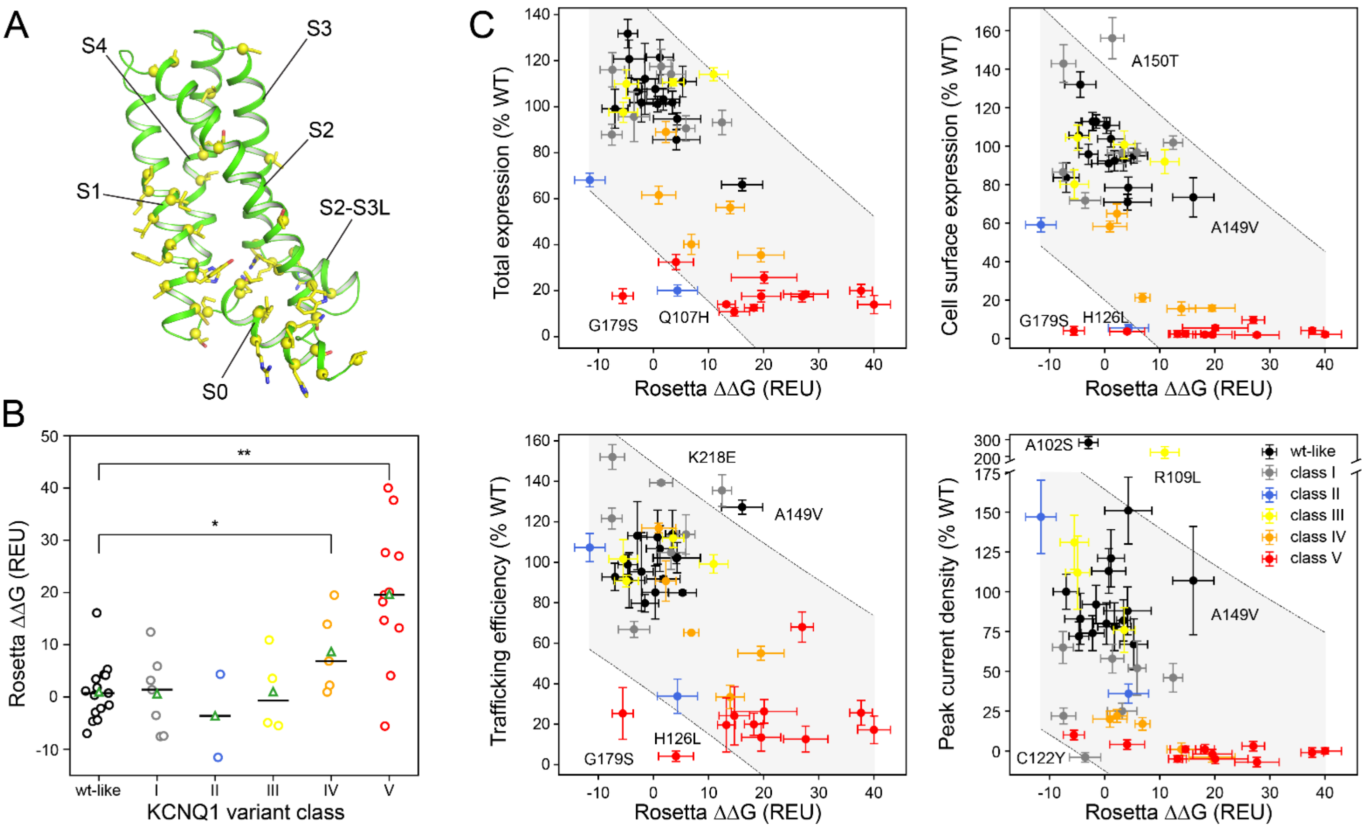

**Figure S7: Predicted stability changes of KCNQ1 VSD variants calculated with the RC model.** (**A**) Location of mutation sites in the KCNQ1 VSD. Backbone sites are mapped by yellow spheres and the native amino acid residue is indicated by yellow sticks. (**B**) Distribution of Rosetta ΔΔG values for the six functionally distinct classes of KCNQ1 VSD variants (i.e. wt-like and classes I to V for non-functional variants) calculated with the RC homology model. The median and average value are drawn as black horizontal line and green triangle, respectively. The median of class IV and V is compared to wt-like variants using a Kruskal-Wallis H-test (* $p < 0.05$, ** $p < 0.01$, *** $p < 0.001$, $n_{\text{wt-like}} = 15$, $n_{IV} = 5$, $n_V = 11$). ΔΔG values for mutations to proline are off-scale ($\Delta\Delta G_{L114P} = 77.1 \pm 4.0$ REU, $\Delta\Delta G_{L131P} = 61.5 \pm 2.5$ REU, $\Delta\Delta G_{L134P} = 63.7 \pm 2.6$ REU, $\Delta\Delta G_{R195P} = 52.7 \pm 1.1$ REU, $\Delta\Delta G_{Q234P} = 82.1 \pm 3.8$ REU, $\Delta\Delta G_{L236P} = 60.3 \pm 4.8$ REU) due to incompatible backbone torsions in the starting model yielding bad backbone and proline ring geometries and were not used in the analysis. (**C**) Correlation plots of total expression level, cell surface expression, trafficking efficiency and channel peak current density versus calculated Rosetta ΔΔG values (mean ± S.E.M.). KCNQ1 variant classes are indicated with different colors. Variants that fall outside or are close to the boundary of the 95% confidence interval for a linear regression model (gray shaded area) are labeled and their structural models are shown in **Figure S8**. The experimental data are from reference (1) and are listed together with the computed ΔΔG values for the RC and AO models in **Table S2**.

Figure S8: Comparison of Rosetta ΔΔG values for KCNQ1 variants calculated with the RC and AO model

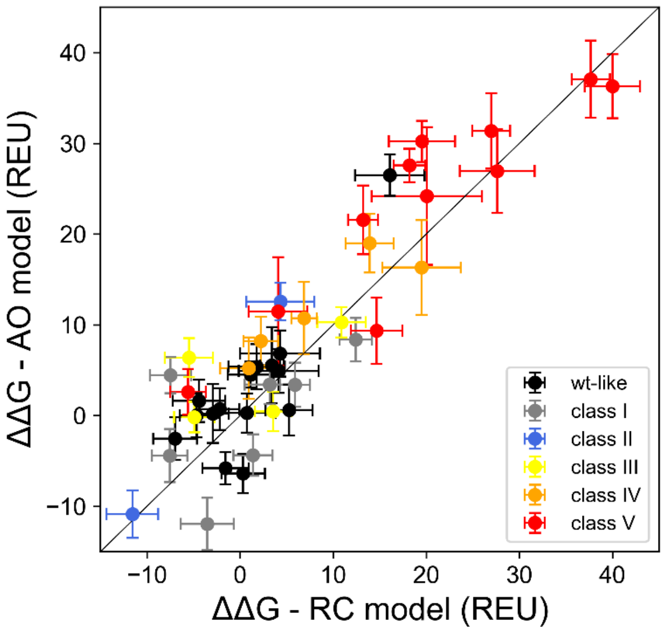

**Figure S8: Comparison of Rosetta ΔΔG values (mean ± S.E.M.) for KCNQ1 variants calculated with the RC and AO model.**

Figure S9: Structural models of selected KCNQ1 variants

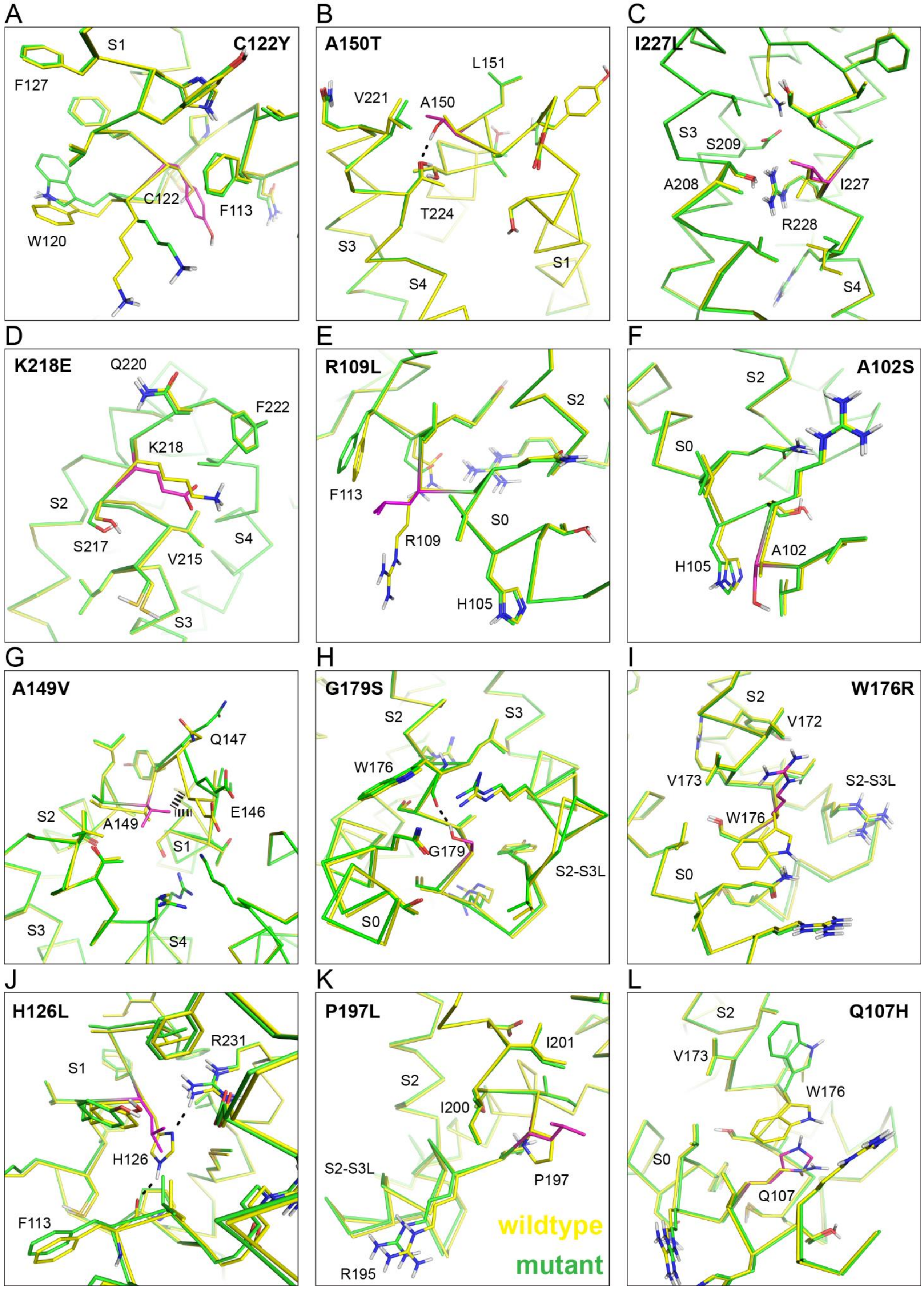

**Figure S9: Structural models of selected KCNQ1 variants.** For each variant, the protein backbone is drawn as ribbon and the side-chains of amino acids within a 5 Å radius of the mutation site are drawn as sticks. The substituted amino acid residue is colored magenta. (**A**) C122Y: has slightly lower expression and trafficking efficiency than wildtype but the channel is not conductive. This is not reflected by the negative ΔΔG (-3.5 REU for RC model and -11.9 REU for AO model) which would indicate a stabilizing effect. (**B**) A150T: has increased cell surface expression and trafficking levels but a current density that is only half of that of wildtype although the protein is predicted to be as stable or more stabilized than wildtype (ΔΔG of 1.4 REU for RC model and -4.4 REU for AO model). Stabilization in the AO model arises from a sidechain-sidechain hydrogen bond with T224 in S4. (**C**) I227: has a negative or moderately increased ΔΔG value (-7.5 REU for RC model and 4.4 REU for AO model) and increased cell surface expression and trafficking which contrasts is very low current density. (**D**) K218E: has an increased positive ΔΔG value (12.8 REU and 8.4 REU for RC and AO model, respectively) (i.e. is predicted to be destabilized) and a current density that is half of that of wildtype but normal cell surface expression and increased trafficking. (**E**) R109L: was predicted to be destabilized (i.e. positive ΔΔG of 10.9 REU and 10.3 REU for RC and AO model, respectively), but has normal expression levels and a current density twofold higher than wildtype. (**F**) A102S: has negative or neutral stability change (ΔΔG of -2.9 REU and 0.2 REU for the RC and AO model, respectively) but a current density almost three times as high as wildtype. (**G**) A149V: belongs to wt-like class with normal or slightly reduced expression levels, but was predicted a high positive ΔΔG (16.1 REU and 26.5 REU for RC and AO model, respectively) which can be explained by steric repulsion between the Val side-chain and the nearby protein backbone in the mutant model (dashed lines). (**H**) G179S: belongs to class V (i.e. severely dysfunctional KCNQ1 variants) but was predicted a low ΔΔG (-5.6 REU and 2.6 REU for RC and AO model, respectively) which is caused by a stabilizing hydrogen bond between the serine side-chain and the W176 backbone oxygen (dashed line). (**I**) W176R: also belongs to class V but the predicted energy loss was only moderate (14.7 REU for RC model and 9.3 REU for AO model) compared to other class V variants. (**J**) H126L: has mildly reduced total expression levels but a strong trafficking defect albeit a moderately increased Rosetta energy (ΔΔG of 4.1 REU and 11.5 REU for RC and AO model, respectively). (**K**) P197L: has reduced total and cell surface expression but was predicted to be more stable than wildtype (ΔΔG of -11.6 REU and -10.9 REU for the RC and AO model respectively). This could indicate a bias favoring leucine over other hydrophobic amino acid residues in the inner and outer hydrophobic layer of the membrane in the RosettaMembrane score function (4). (**L**) Q107H: was previously assigned class II but may be re-assigned class V based on its low expression/trafficking levels, low conductance and positive ΔΔG (4.3 REU and 12.6 for RC and AO model, respectively).

Figure S10: Mapping of experimental KCNE1 contact sites onto KCNQ1 models

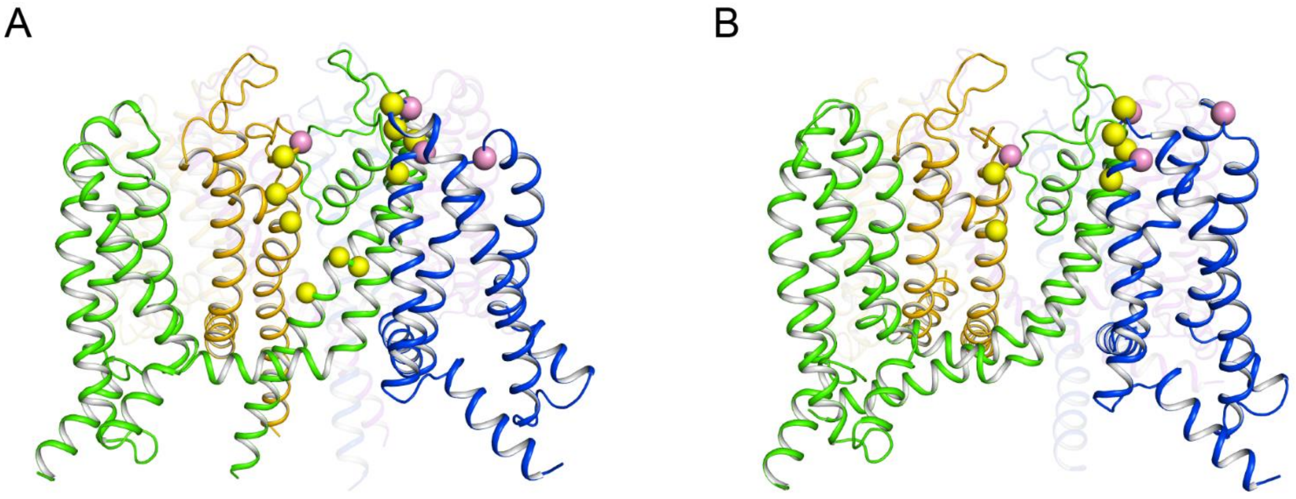

**Figure S10: Mapping of experimental KCNE1 contact sites onto KCNQ1 models.** Cartoon representation of (**A**) the RC and (**B**) the AO model. Chain A (green), the VSD of chain B (blue) and the PD of chain D (orange) are shown in the front. KCNE1 interaction sites are represented as spheres. The subunits in the back are drawn transparent for clarity. Experimental restraints are from references (5-11). KCNQ1 positions with restraints specific for the RC or AO state, respectively, are illustrated as yellow spheres. Restraints for which the channel state was unclear are indicated as pink spheres.

### Supporting References

1. Huang H, Kuenze G, Smith JA, Taylor KC, Duran AM, Hadziselimovic A, et al. Mechanisms of KCNQ1 channel dysfunction in long QT syndrome involving voltage sensor domain mutations. Sci Adv. 2018;4(3):eaar2631.
2. Ortiz AR, Strauss CEM, Olmea O. MAMMOTH (Matching molecular models obtained from theory): An automated method for model comparison. Protein Sci. 2002;11:2606-11.
3. Larkin MA, Blackshields G, Brown NP, Chenna R, McGettigan PA, McWilliam H, et al. Clustal W and Clustal X version 2.0. Bioinformatics. 2007;23(21):2947-8.
4. Duran AM, Meiler J. Computational design of membrane proteins using RosettaMembrane. Protein Sci. 2018;27(1):341-55.
5. Chan PJ, Osteen JD, Xiong D, Bohnen MS, Doshi D, Sampson KJ, et al. Characterization of KCNQ1 atrial fibrillation mutations reveals distinct dependence on KCNE1. J Gen Physiol. 2012;139:135-44.
6. Tapper AR, George AL. Location and orientation of minK within the I-Ks potassium channel complex. Journal of Biological Chemistry. 2001;276(41):38249-54.
7. Chung DY, Chan PJ, Bankston JR, Yang L, Liu G, Marx SO, et al. Location of KCNE1 relative to KCNQ1 in the I(KS) potassium channel by disulfide cross-linking of substituted cysteines. Proc Natl Acad Sci U S A. 2009;106:743-8.
8. Li P, Liu H, Lai C, Sun P, Zeng W, Wu F, et al. Differential modulations of KCNQ1 by auxiliary proteins KCNE1 and KCNE2. Sci Rep. 2014;4:4973.
9. Strutz-Seebohm N, Pusch M, Wolf S, Stoll R, Tapken D, Gerwert K, et al. Structural basis of slow activation gating in the cardiac I Ks channel complex. Cell Physiol Biochem. 2011;27:443-52.
10. Xu X, Jiang M, Hsu K-L, Zhang M, Tseng G-N. KCNQ1 and KCNE1 in the IKs channel complex make state-dependent contacts in their extracellular domains. J Gen Physiol. 2008;131:589-603.
11. Wang YH, Jiang M, Xu XL, Hsu K-L, Zhang M, Tseng G-N. Gating-related molecular motions in the extracellular domain of the IKs channel: implications for IKs channelopathy. J Membr Biol. 2011;239:137-56.
